# Floral phenotypic divergence and genomic insights in an *Ophrys* orchid: Unraveling early speciation processes

**DOI:** 10.1101/2024.03.21.586062

**Authors:** Anaïs Gibert, Bertrand Schatz, Roselyne Buscail, Dominique Nguyen, Michel Baguette, Nicolas Barthes, Joris A.M. Bertrand

## Abstract

- Adaptive radiation in *Ophrys* orchids leads to complex floral phenotypes that vary in scent, color and shape.
- Using a novel pipeline to quantify these phenotypes, we investigated trait divergence at early stages of speciation in six populations of *Ophrys aveyronensis* experiencing recent allopatry. By integrating different genetic/genomic techniques, we investigated: (i) variation and integration of floral components (scent, color and shape), (ii) phenotypes and genomic regions under divergent selection, and (iii) the genomic bases of trait variation.
- We identified a large genomic island of divergence, associated with phenotypic variation in particular in floral odor. We detected potential divergent selection on macular color, while convergent selection was suspected on floral morphology and for several volatile olfactive compounds. We also identify candidate genes involved in anthocyanin and in steroid biosynthesis pathways associated with standing genetic variation in color and odor.
- This study sheds light on early differentiation in *Ophrys*, revealing patterns that often become invisible over time, i.e., the geographic mosaic of traits under selection and the early appearance of strong genomic divergence. It also supports a crucial genomic region for future investigation and highlights the value of a multifaceted approach in unraveling speciation within taxa with large genomes.

## Introduction

How new species arise remains a fundamental question in evolutionary biology. The genomic era has deepened insights into the connections between genomic architecture, genetics, phenotypic divergence, and reproductive isolation. Nevertheless, our understanding about the link between the emergence of divergent phenotypes and the build-up of genetic changes that cause reproductive isolation remains limited (Schlüter & Rieseberg, 2022). The genus *Ophrys*, or bee orchids, provides a compelling model for studying the early stages of speciation. Firstly, it represents a very recent and intense adaptive radiation, with one of the highest speciation rates recorded so far among angiosperms (Breitkopf *et al*., 2015), and numerous species still at the onset of divergence (e.g., Gibert *et al*., 2023, ca. 1500 generations). Second, the diversification within *Ophrys* is thought to be due to a remarkable pollination syndrome called sexual deception (Baguette *et al*., 2020). Here, the key innovation is for the flowers to mimic female insects using visual, tactile and olfactory cues and are pollinated by male insects that mistake the flower for a female during a “pseudocopulation”. As a result, the *Ophrys* model exhibits a multifaceted floral phenotype, including variations in color, shape and odor (Schiestl, 2005; Schiestl & Schlüter, 2009), which may contribute to reproductive isolation as an immediate by-product (the so-called ‘magic traits’, Schlüter, 2018).

Extensive research has been conducted on the *Ophrys* model to determine the significance of floral characteristics in speciation. The findings demonstrate the leading role of scent characteristics in all *Ophrys* species investigated in attracting pollinators, eliciting copulation and ensuring reproductive isolation (Ayasse *et al*., 2000, 2003; Schiestl *et al*., 2000; Schiestl & Ayasse, 2002; Mant *et al*., 2005; Schiestl, 2005; Vereecken & Schiestl, 2009; Gervasi *et al*., 2017). However, the *Ophrys* model has received little attention in exploring the genomic bases of speciation to date, at the notable exception of the candidate gene approach developed by Schlüter, Xu and collaborators (Schlüter *et al*., 2011; Xu *et al*., 2012, 2017). This is partly due to its large genome size (approximately 5-7 Gb for diploid species, Russo *et al*. 2023) and also to the challenge of obtaining quantitative data from all aspects of this complex phenotype. Recent advances in the extraction of quantitative data from *Ophrys* pictures (e.g., Gibert *et al*., 2022) and the publication of a reference genome in *Ophrys* (Russo *et al*. 2023) present new perspectives for investigating speciation in this system.

In this study, our focus lies on three core areas: i) exploring the patterns of variation and integration of traits within the complex phenotype both between and within populations, ii) elucidating whether these traits are under divergent or convergent selection, and iii) unraveling their genetic architecture. Here, we build on the hypothesis formalized by Baguette *et al*. (2020), which suggests that pollinator shifts in *Ophrys* result from divergent and variable olfactory traits that lead subsequently to directional divergence in color and shape to enhance attractiveness to new pollinators. This is why we expect to observe divergent selection on some olfactory compounds, possibly followed by divergent selection on color and morphological aspects (depending on the stage of the speciation process). Furthermore, we anticipate significant variation in scent within and between populations, as well as some integration with color traits resulting from shared biological pathways identified in other plants. Additionally, adhering to Berg’s theory (1960), we predict reduced variance in floral morphological traits when compared to overall plant traits. Finally, we expect to detect variations in genes related to the biosynthetic pathways of volatile organic compounds (VOCs) such as the genes coding for stearoyl-ACP desaturase (SAD) enzymes and which induce the main odor differences between *Ophrys sphegodes* and *O. exaltata* (Schlüter *et al*., 2011; Xu *et al*., 2012, 2017).

Here, we study the process of diversification in *Ophrys aveyronensis*; a system with two allopatric taxa at an early stage of speciation (*Ophrys aveyronensis* subsp. *aveyronensis* in France and *Ophrys aveyronensis* subsp. *vitorica* in Spain). While no pollinator shift has yet been reported in this system, patterns of recent genetic divergence (Gibert *et al*., 2023) and cryptic phenotypic differentiation in morphology and color - but never in odor - have been reported (Gibert *et al*., 2022). Whether these differences are adaptive or simply due to genetic drift has never been investigated. In this study, we used a RADseq-like protocol to generate data and recorded morphological, color and olfactory traits for 86 individuals in six populations using a recently published pipeline for extracting quantitative data from images (Gibert *et al*., 2022). We combined i) variance and network analyses of traits to investigate the pattern of variation and the relationship between odor, color and morphological traits, ii) a quantitative genetic approach (*P*ST-*F*ST approach) to identify adaptive traits under selection, iii) reverse genetic approaches to identify regions of the genome with signatures of divergent selection (genome scan) and/or associations between genotypes and environmental factors (GEA), and iv) a forward genetic approach (GWAS) to understand the underlying genetic architecture of trait variation. We used the recently sequenced, assembled and annotated genome of a closely related species, *O. sphegodes*, to determine the potential functional role of the identified SNP outliers.

## Material and Methods

### 1. Study system

*Ophrys sphegodes* Mill. subsp. *aveyronensis* (Wood, 1983), also defined as *Ophrys aveyronensis* (Delforge, 2016), was first described as endemic to the Grands Causses region (southern France). Later, phenotypically similar populations were sporadically discovered in Northern Spain (Hermosilla & Sabando, 1998; Hermosilla & Soca, 1999; Benito Ayuso, 2019). While no clear differences in ecology and pollinator (a solitary bee: *Andrena hattorfiana*) were found between the French and Iberian populations (Paulus, 2017), these populations have been shown to be diverging, suggesting an early stage of speciation (Gibert *et al*., 2022; 2023) and are currently considered as subspecies: *O. a.* subsp. *aveyronensis* (in France) and *O. a.* subsp. *vitorica* (in Spain).

### 2. Collection sites and plant material

In June 2019, we collected samples from six populations representative of the disjunct geographic distribution of the *Ophrys aveyronensis* species complex: three populations in south of France (Guilhaumard, Lapanouse-de-Cernon and Saint- Affrique) and three populations in north of Spain (Valgañón, Larraona and Bercedo). From each population, between 13 and 15 individuals were genotyped, measured, and photographed for a total of 86 individuals. These individuals are the same than the one investigated in Gibert *et al*., (2022, 2023). A permit (n°2019-s-16) was obtained from the DREAL of Occitanie (Direction Régionale de l’Environnement et de l’Aménagement du Territoire) to collect samples in France, where *Ophrys aveyronensis* is legally protected. No permit was required in Spain.

Each population was characterized by a set of 19 bioclimatic variables extracted from the WORLDCLIM database (version 2, Hijmans *et al*., 2005). These bioclimatic variables represent annual trends, extremes and limits of climatic factors related to temperatures and precipitations over the period from 1970 to 2000. To reduce the multidimensional bioclimatic dataset to a few uncorrelated factors, we performed a Principal Component Analysis using the R package vegan (Oksanen *et al*., 2009), and selected the two-firsts PCs of the PCA as a synthetic climatic data in the analyses. These two-firsts PC axes of the temperature-precipitation PCA were retained and accounted for 70% of the total variance of the dataset. The strongest correlations with PC1 are annual precipitation, temperature seasonality and precipitation of the wettest month.

### 3. Phenotypic traits

We examined differentiation at 45 phenotypic traits (see boxplots in supplementary Fig S1a:f). Floral and plant-level morphological traits included the width and length of the labellum, sepal and petal, and the between pseudo-eyes distance. Plant-level traits included plant size, stem diameter, distance from ground to first flower, distance from first to second flower, number of flowers, number of buds, and number of buds and flowers. Colors information is provided as eight color variables corresponding to the quantity of pixels in eight bins of colors with similar boundaries (Fig 1). See further details in Gibert *et al*. (2022).

**Figure 1.**
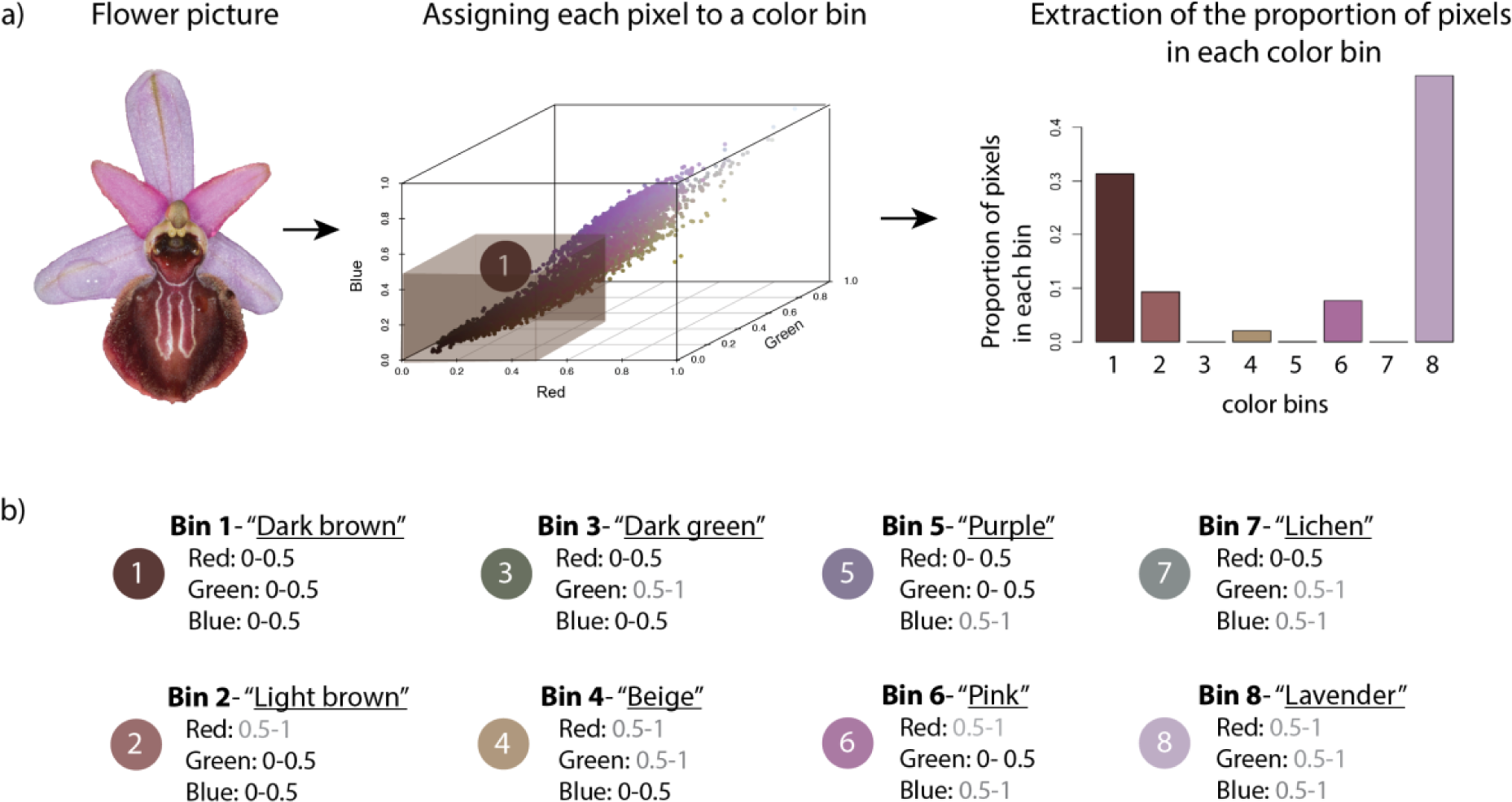
Approach used for extracting quantitative color data in flowers (**a**) main steps using the R package “colordistance” (Weller & Westneat, 2019), (**b**) information on colors bins.

In total 46 individuals were sampled for floral compounds using solid phase microextraction (SPME) and analyzed by GC-MS methods (see Supporting Information Methods S1). This non-destructive technique has already been successfully used to extract floral compounds in *Ophrys* species (Joffard *et al*., 2016).

We identified and analyzed the most abundant compounds (i.e 23 compounds, present in at least 20% of the individuals, and constituting more than 0.5% of the floral bouquet). These 23 compounds make up over 67% of the floral bouquet, with each compound contributing 0.5-9% of the total (Table 1). All analyses were performed on the relative proportions of these 23 compounds, but also on their occurrence (0/1) when it was not found in at least 20% of the individuals.

**Table 1.**
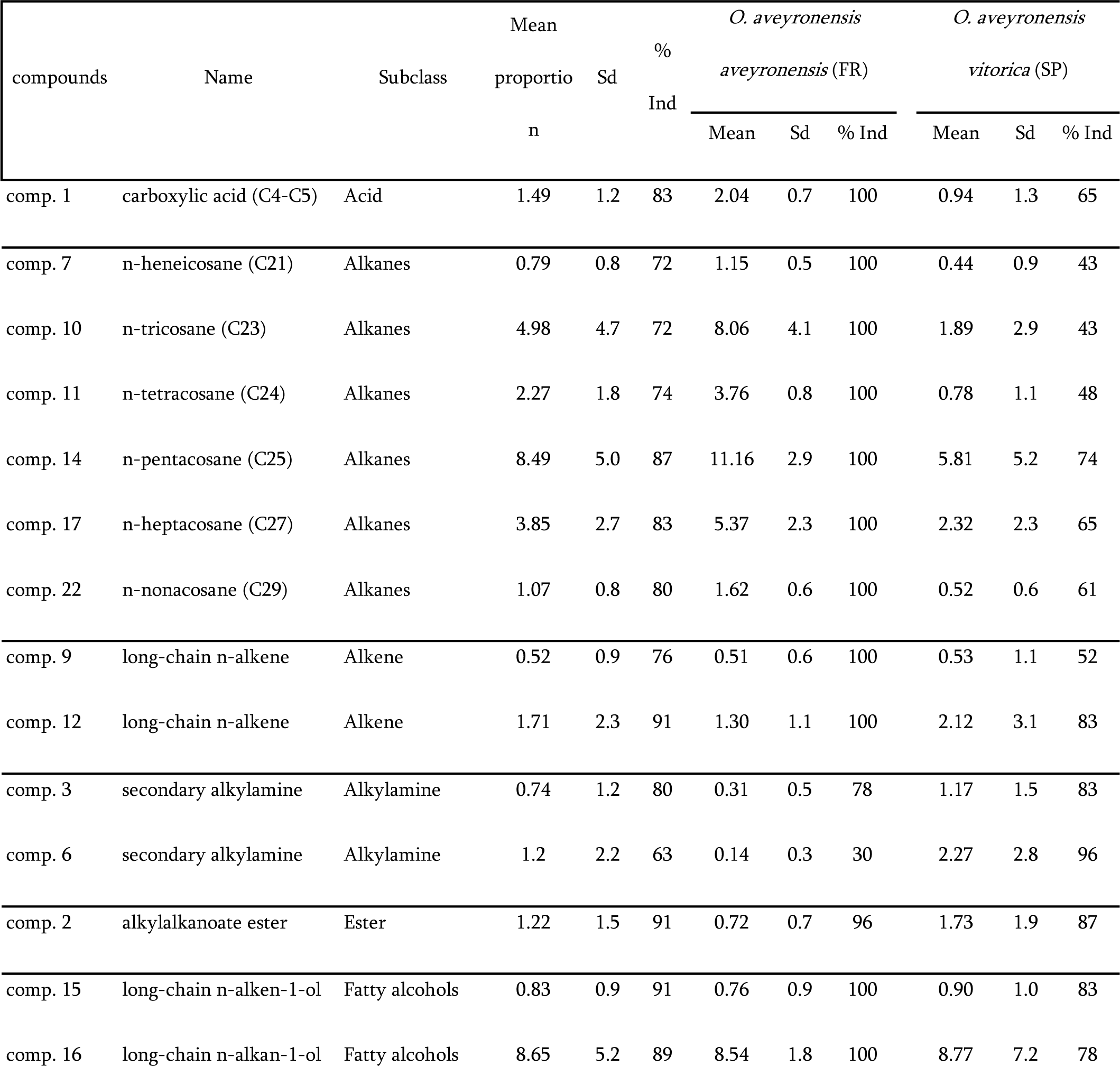
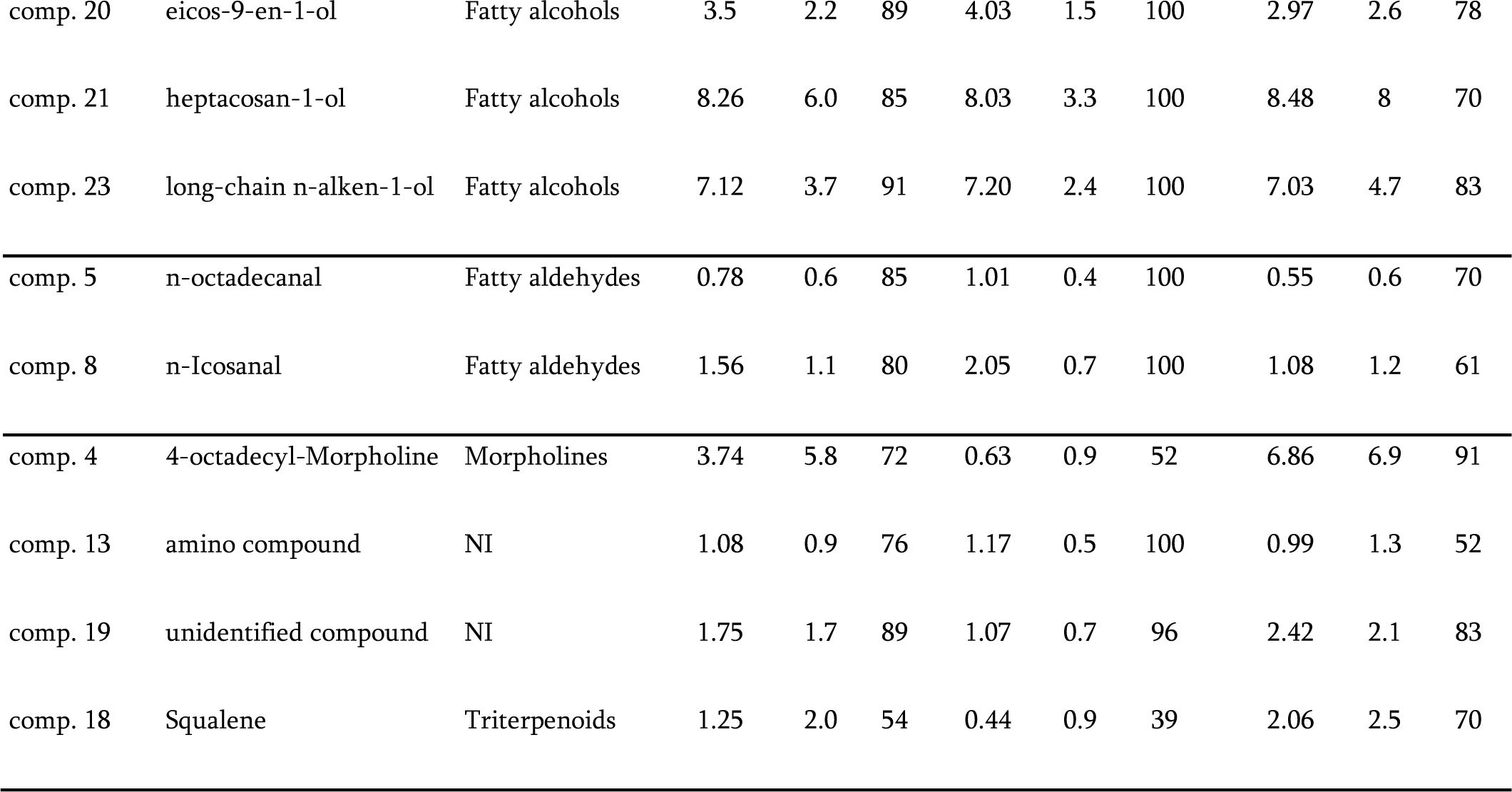
Compounds identified by gas chromatographic analyses on flower labella of the two subspecies of *Ophrys aveyronensis*. Mean: mean relative proportion ±sd. % Ind: percentage of individuals emitting the compound. Compound numbers refer to order by size.

### 4. DNA extraction, genotyping and variant calling

Genomic DNA extraction, library preparation and genotyping were subcontracted to LGC Genomics GmbH (Berlin, Germany). A fractional genome sequencing strategy called normalized Genotyping-by-Sequencing (nGBS) was used to subsample the *Ophrys aveyronensis* genome (see Gibert *et al*. 2022 for details). We used the sequence of the genome of *Ophrys sphegodes* (Russo *et al*. 2023) as a reference to build loci and call SNPs with the pipeline Stacks v.2.60 (Catchen *et al*., 2011; 2013). See additional details in supporting informations (Methods S2).

### 5. Phenotypic differentiation between regions and populations

Here we expect to identify phenotypic variation between regions, but also between populations, since the population is the scale at which natural selection operates.

#### a. Phenotypic differentiation

Differentiation between regions and/or among populations was tested independently on each traits using generalized linear models (GLMs). In each model, fixed effects included the regional effect (France vs. Spain) and the population effect nested within the regional effect. Several traits (reproductive structure, several odor compounds and color bins 3-“Dark green”, 5-“Purple” and 7-“Lichen”, see Fig 1) were log-transformed and/or outliers removed to meet model assumptions (e.g. heteroscedasticity, no over- or under-dispersion, no outliers). All traits were modeled with a Gaussian distribution, except the number of buds which was modeled with a Poisson distribution (with a log link function). The chosen link functions satisfied the assumptions of the models. All GLMs were implemented in R using the lme4 package (Bates *et al*., 2015). Model assumptions (e.g. uniformity, dispersion, outliers) were checked using the R package Darmha (Hartig, 2020). Regional and population effects were only tested for traits that had positive values in more than five individuals in at least two populations (see Supporting Information, Table S1).

#### b. Floral bouquet differentiation

To test whether the floral bouquet overall differs in *Ophrys aveyronensis* the blends of the two taxa were compared between regions and also between populations. We used two approaches: i) sparse partial least squares-discriminant analysis (sPLS-DA) and ii) random forest models (RF). The sPLS-DA were performed using the R package mixOmics (Rohart *et al*., 2017) and it performs variable selection and classification in a one-step procedure (Lê Cao *et al*., 2011). We used cross-validation tests to estimate the classification error rate (M-fold method) and its statistical significance (permutation test with 999 permutations). The RFs with k-fold cross-validation were implemented in R using the randomForest package (Liaw & Wiener, 2015). We ran models with 500 trees, and we selected an optimal value for mtry based on cross-validation performance (the out-of-bag-error estimate, using the tuneRF function in R), and the node size was set to default. We evaluated the accuracy of the classifier as measured of performance, and we reported the out-of-bag-error (OOB error).

### 6. Patterns of traits variation and integration

To compare variation across traits, we calculated trait variation within the populations as a dimensionless coefficient of variation (CV) = σ / μ, with σ being the standard deviation and μ the mean of the individuals sampled. To avoid potential scaling artifacts, we focused exclusively on one dimensional traits (length and count vs. area and volume) and we discarded the traits for which means and standard deviations were uncorrelated (Pélabon *et al*., 2020, see list in Table S1, Criterion 2). We used linear mixed models to test whether some trait categories (floral morphology, plant level morphology, color and odor traits) were more variable than others. Fixed effects included trait category and regions (France vs. Spain). Random effects included specific traits. Data were log-transformed to meet the assumptions of the model. We report model-averaged parameter estimates derived from multi-model inference (Grueber *et al*., 2011; Harrison *et al*., 2018). This approach allowed us to incorporate uncertainty by weighting the parameter estimate of a model by that model’s Akaike weight. Model averaging was performed in R using the package MuMIn (Barton, 2009). Model assumptions (e.g. uniformity, dispersion, outliers) were checked using the R-package Darmha (Hartig, 2020) with R.3.5.0 software (R Core Team, 2018).

We also analyzed the spatial structuring of the variance in color, odor, floral morphology and plant-level morphology. Percentages of explained variance between regions and between populations within regions were computed as individual adjusted R2 using Ezekiel’s formula through variation and hierarchical partitioning with the R package “rdacca.hp” (Lai *et al*., 2022).

Relationships between plant traits were visualized using a plant trait network (PTN), where traits were set as nodes and trait-trait relationships were set as edges following the approach of He *et al*. (2020). We also identified modules of traits that are densely connected to each other but sparsely connected to other modules in this network (Newman & Girvan, 2004; Newman, 2006). See more details in supporting information (Methods S3). Traits within a cluster would be studied concomitantly in subsequent GWAS analysis.

### 7. Quantitative genetic approach: *P*ST-*F*ST approach

To detect putative selection signals, we compared phenotypic differentiation for each trait (*P*ST, Leinonen *et al*., 2008) with neutral genetic differentiation (*F*ST, Weir & Cockerham, 1984) between populations. This genotype-phenotype comparison has three possible outcomes: (i) *P*ST = *F*ST, indicating that the level of phenotypic differentiation is equal to the neutral genetic differentiation between populations, a pattern consistent with neutral processes (e.g. genetic drift etc.) as the main mechanisms explaining phenotypic divergence. (ii) *P*ST > *F*ST, indicating that the degree of phenotypic differentiation is greater than expected under the assumption of neutrality (no selective pressure). This pattern suggests that directional or divergent selection may be involved in promoting phenotypic divergence between populations. (iii) *P*ST < *F*ST, indicating that the degree of phenotypic differentiation is less than expected under the assumption of neutrality (no selective pressure). This pattern suggests that stabilizing selection and/or canalization might limit phenotypic divergence between populations. *P*ST were calculated using the equation proposed by Brommer (2011), and using a Bayesian generalized linear mixed model (Bertrand *et al*., 2016; Seeholzer & Brumfield, 2018). See more details in supporting information (Methods S4).

### 8. Forward genetic approaches

We used two forward genetic approaches: genome scan (GS) and genome-environment association (GEA). If a genomic region of interest is identified with these methods, we would infer the population structure observed with and without this region using a Principal Component Analysis (PCA) coded as a genlight object with the package adegenet (Jombart & Ahmed, 2011).

Genome scan approaches allowed to detect a signature of divergent selection without assuming the effect of any particular trait or environmental variable, allowing to investigate the existence of highly differentiated SNPs across the genome. To do this, we used three population differentiation methods: pcadapt, snmf and baypass. The null assumption tested is that each SNP is not under selection and therefore not overly differentiated from the genomic background. The default values of the three programs were used for all their internal parameters. See details on the implementation of each method in Appendix 1 - Methods. For pcadapt, to identify outliers associated with population divergence we ran the function with K=5 (optimal number of genetic clusters inferred), but we also ran the function with K=2 (subdividing the two taxa) to identify outliers for regional divergence.

Genotype-environment association (GEA) approaches detect SNPs that are significantly associated with environmental variables. The null hypothesis is no correlation between SNP allele frequencies and the environmental variable; genetic variation is only due to genetic drift and isolation. Here, we used the latent factor mixed model (LFMM), a mixed linear model, to test for associations between SNP allele frequencies and two synthetic bioclimatic variables (PC1 and PC2), while correcting for neutral population structure through the latent factor (here K=5 based on previous results, Gibert *et al*., 2023). We used the implementation of LFMM in the R package LEA version 3.6.0 (lfmm2 function, Frichot & François, 2015) to perform the analysis. See more details in supporting information (Methods S5).

### 9. Reverse genetic approaches: Genotype-phenotype associations

As there is no best model for all traits, we ran two genome-wide association studies (GWAS) models (FarmCPU: fixed and random model circulating probability unification, and BLINK: Bayesian information and Linkage disequilibrium Iteratively Nested Keyway) and a latent factor model (LFMM) to detect outlier loci potentially affected by divergent selection on phenotypic traits. All models, control for population structure. LFMM is performed as for the GEA approach but with phenotypic traits as independent variables, using the LEA package (Frichot & François, 2015). FarmCPU is a hybrid model that combines the advantages of mixed linear models and stepwise regression (Liu *et al*., 2016). BLINK is based on fixed effects models incorporating Bayesian information criteria (Huang *et al*., 2019). Both models were implemented in the R package GAPIT (Wang & Zhang, 2021). P-values are adjusted as implemented in GAPIT, following a false discovery rate (FDR) control procedure (Benjamini & Hochberg, 1995). Quantile–quantile (QQ) plots were used to assess the extent of accordance between the observed and expected p-values. See more details in supporting information (Methods S6).

For traits forming clusters in the plant trait network, we performed a PCA for each cluster, and performed GWAS (with lfmm2, FarmCPU and BLINK) on the top two PC axes and on a combined axis.

### 10. Annotation of candidate SNPs

For each SNP, we searched for CDS located 5-kb upstream and downstream from the candidate outliers using *Ophrys sphegodes* as reference. The function of these CDS was identified using diamond blastx against a UniProt database restricted to the Orchidaceae family (The UniProt Consortium, 2023) with e-values ≤ 0.01 and bitscore ≥ 100, ii) annotation of the region of interest done by Russo *et al*. (2023) and iii) annotation via EGGNOG-MAPPER (Cantalapiedra *et al*., 2021) with an alignment p-values ≤ 0.001 and an alignment score ≥ 60.

When a region of genomic divergence was found, we performed gene ontology (GO) enrichment analysis using the topGO package (parameters: Fisher’s exact test. weight01 algorithm, minimum node size of 3; v.2.26.0; Alexa & Rahnenfuhrer, 2023). The function (GO) of filtered predicted coding genesis annotated with the EGGNOG-MAPPER version 2.0.1 (Cantalapiedra *et al*., 2021) using CDS for these genes, with an alignment p-values ≤ 0.001 and an alignment score ≥ 60. We contrasted them with all the annotated protein-coding genes of *O. sphegodes*. Results were considered significant at p-values < 0.01. Corrections for multiple testing were not performed, as recommended in the topGO manual.

## Results

### Differentiation at the regional and population level varied according to the trait considered

The majority of phenotypic traits showed a significant phenotypic differentiation between regions and/or populations (32/45 traits, Table S2). A notable exception was found for most floral traits (petal width, labellum length, petal length, sepal length), few volatile olfactive compounds (VOCs, two alkenes: comp. 9 and 12, four fatty alcohols : comp. 15, 16, 21, 23; and one alkylamine : comp. 3 as occurrence) and one color trait (proportion of pixels in the bin of color 2- “light brown”), where no evidence of differentiation at any geographical scale was identified.

Overall, the floral bouquet of *Ophrys aveyronensis* differs between regions and populations; similar VOCs but distinct ratios of the same compounds were emitted (Table 1). The blends were well differentiated between regions as shown by the non-overlapping scatterplot, the heatmap (Fig 2 a, b and c), and by the cross-validation tests (classification error rate < 0.05, area under the curve (AUC) > 0.98 with a p-value < 3.10-8). The random forest model showed a 100% accuracy, and a OOB error rate of 5.2% between subspecies. In contrast, to distinguish between the six populations based on their blends was not efficient (Fig S5); the RF between populations showed a 42.86% accuracy, and a OOB error rate of 60.53%.

**Figure 2.**
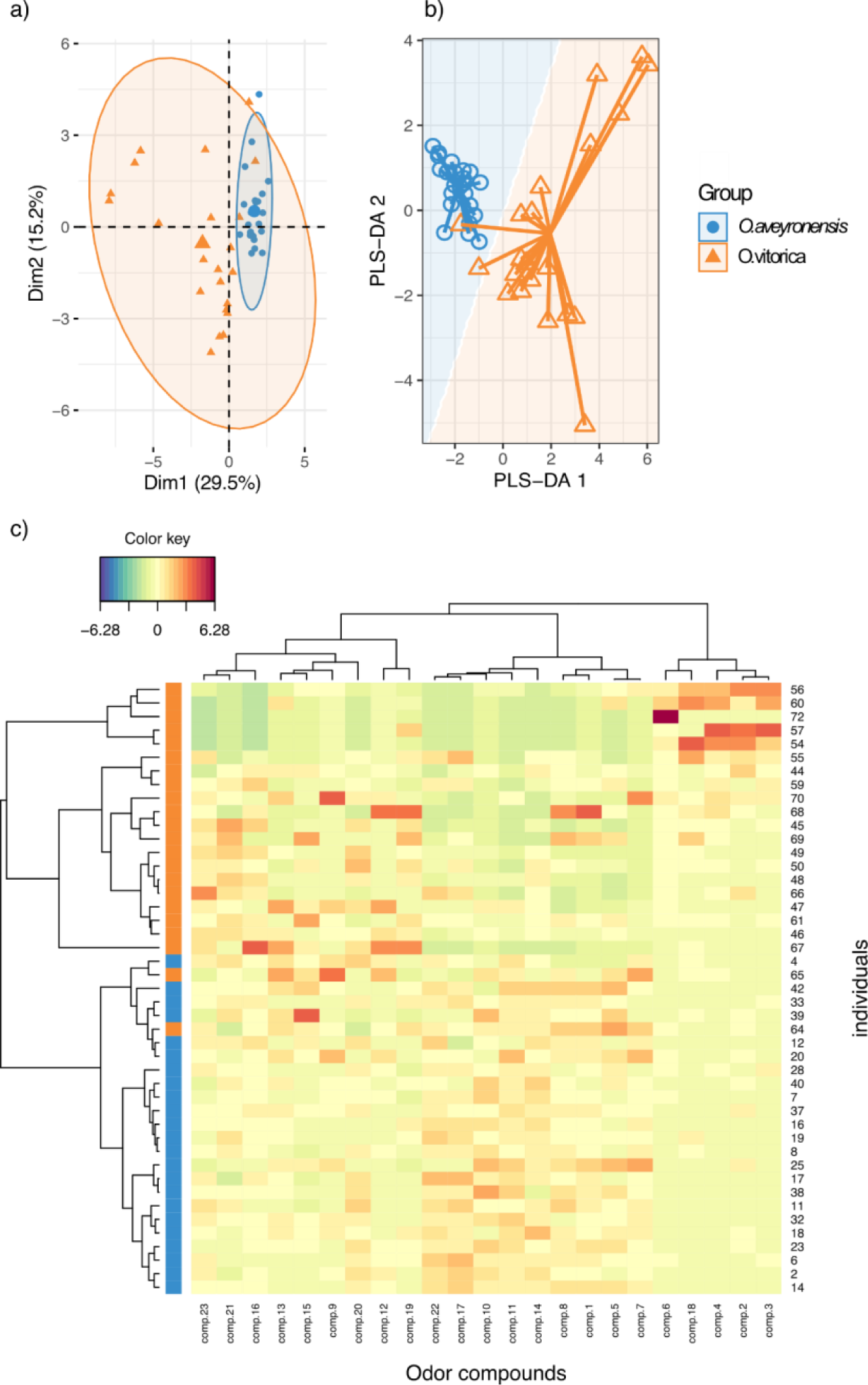
Differentiation of floral bouquet between subspecies with supervised analysis. (**a**) PCA, (**b**) sPLS-DA sample plot with confidence ellipse plots, (**c**) Clustered Image Map (Euclidean Distance) with the odor compounds in columns, and the individuals in rows.

### Variation in odor and color traits, but no variation in floral morphology

Overall, floral morphological traits were significantly less variable than other traits (Fig 3 a and b). By contrast, odor traits and color traits were the most variable. For all trait categories, most of the variation occurred within the population (i.e. between individuals, 75 to 97% of the variance, Fig 3 c). Significant variation between regions and/or populations within regions occurred for odor traits (region: 12%, p-value = 0.001, populations: 10%, p-value = 0.009), color traits (populations: 24.5%, p-value= 0.01) and plant-level morphological traits (region: 2.5%, p-value = 0.022, populations: 11.6%, p-value = 0.001). Floral morphological traits showed no significant variation at any scale (regions: 0.9%, p-value= 0.111, populations: 2.5%, p-value= 0.067).

**Figure 3.**
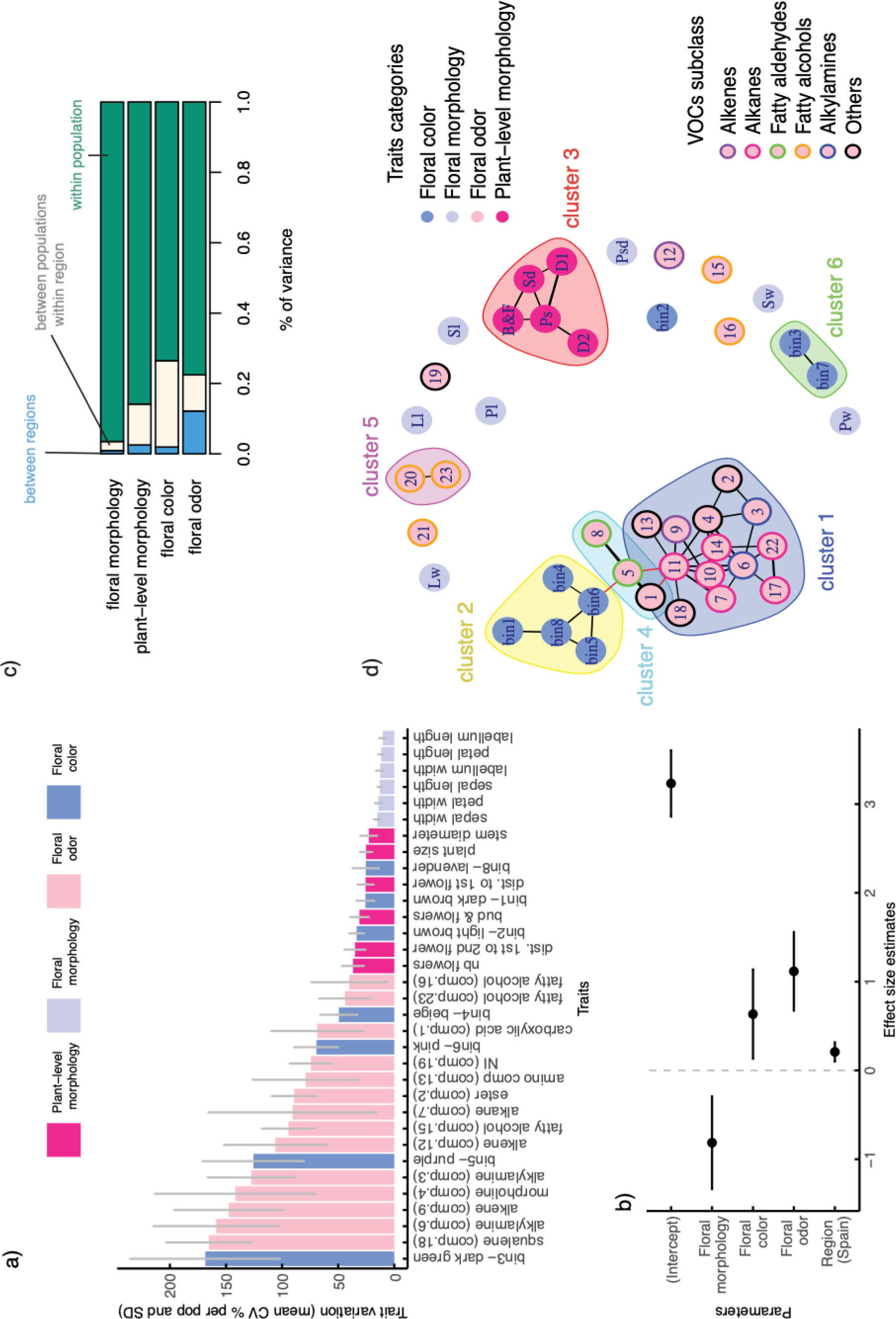
Patterns of trait variation and co-variation. (**a**) Histogram showing variation by traits (mean CV by population ±SD across populations), (**b**) Effect-size plot on CV from an averaged linear mixed model with trait categories and regions as fixed effects, and populations and traits as random effects. Effect sizes are reported with 95% confidence intervals (CI) and are significant if the CI does not overlap with zero. R2 = 0.78, and R2-adjusted = 0.84. (**c**) Spatial variance partitioning between regions (blue), between populations within region (white) and within populations (i.e. individual scale in green), (**d**) Trait correlation network. Only significant correlations (p-values < 0.05) with strength equal to or greater than 0.5 are shown. Ps: plant size; Sd: stem diameter; D1: distance to first flower; D2: distance from first to second flower; B&F: number of buds and flowers; Sl: sepal length; Sw: sepal width; Pl: petal length; Pw: petal width; Ll: label length; Lw: label width; Psd: pseudo-eye distance. From n°1 to 23: odor compounds; see table 1 for detailed name of compounds.

### Independence of floral morphological traits, clustering and relationship of odor and color traits

The trait network suggests that most traits are likely to be connected as modules (Fig 3 d). One interesting cluster is composed of all alkanes (heneicosane, tricosane, tetracosane, pentacosane, hetpacosane, nonacosane) and one on the two alkenes identified in the floral bouquet (comp. 9, cluster n°1, Fig 3d). All floral morphological traits formed independent nodes, as well as few VOCs (one alkene: comp. 12, and several fatty alcohols: comp. 15, 16, 21) and one color traits (bin 2 -“light brown”). These traits have been identified as non-variants across population and region in the previous analysis. Modularity score was maximal for this clustering (Q = 0.57 vs Q = 0.41 for a clustering done with trait categories). This trait-network highlighted some relationship between clusters, in particular a relationship between an odor and a color cluster.

### Potential signatures of divergent selection on color traits and convergent selection on floral and odor traits

Three color traits (color bins 3 “Dark green”, 5 “Purple” and 7 “Lichen”) showed *P*ST > *F*ST (at c/h2=1) with non-overlapping CIs, suggesting divergent selection (Fig 4, Fig S6 a & b). When the robustness of these conclusions was examined with respect to variation in c and h2, we found low values of critical c/h2 (< 0.2) for bin 3 and bin 7, but not for bin 5. The critical c/h2 for bin 5 was 0.7, which casts doubt on the robustness of inference regarding the putative effect of selection on the observed divergence in this trait.

**Figure 4.**
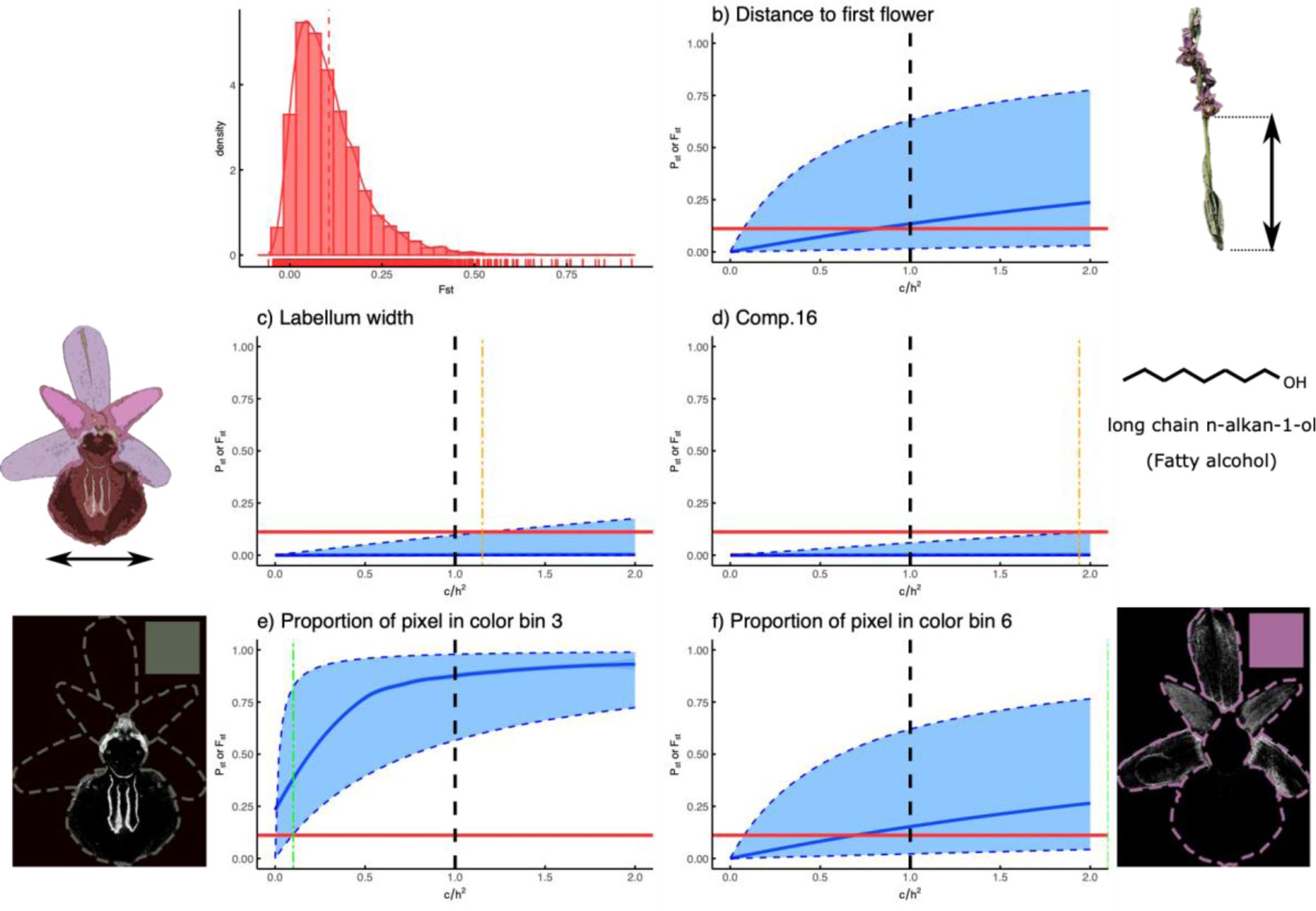
Plot (**a**) is the distribution of *F*ST over loci (mean= 0.11, CI 95% = 0.10- 0.112). Plots (**b**, **c**, **d**, **e**, **f)** comparisons of phenotypic differentiation (*P*ST) with neutral genetic differentiation (*F*ST) for several traits. *P*ST (blue line) is plotted with CIs (dotted blue line) as a function of c/h2.

Three odor traits, all fatty alcohols (comp. 16, comp. 21, comp. 23), showed *P*ST < *F*ST (at c/h2=1) with non-overlapping CIs, suggesting convergent selection (Fig 4 d, Fig S6 c). We found high values of critical c/h2 (above 1) suggesting fairly robust inferences of convergent selection (or constraint). For several floral traits (petal width, sepal width, labellum length and width), *P*ST tended to be lower than neutral expectation at c/h2=1 (Fig 4 and S6 a & b), suggesting convergent selection (or canalization). Yet, estimates of variance components had low precision when *P*ST were close to zero, and CIs overlapped *F*ST for several of these traits. Only labellum width and petal width exhibited non overlapping CIs at c/h2=1, and high critical c/h2 values (above 1) suggesting convergent selection (or canalization).

Several traits were excluded from this analysis; those for which the models did not converge and/or did not fulfill the model assumptions (Table S1, criterion 3). Thus, for petal and sepal length, for most alkanes (5/9, comp. 10, 11, 14, 17, 22), one alkene (comp. 12) and several other volatile olfactory compounds, the diagnostics indicated that the models did not converge correctly. For any other traits tested, there was no evidence that the observed differentiation (or convergence) differed from the assumption of neutrality (no selective pressure).

### A large region of genomic divergence on chromosome 2 identified by forward genetic approaches

The genome scan methods detected a divergent region of ca. 7 Mb at the end of chromosome 2 (pos: 336910807 - 344195882, Fig 5). This region was not detected by GS methods focusing on subspecies divergence (snmf and pcadapt with K=2) suggesting that this divergence occurred rather at population level. This is confirmed by the study of population genetic structure with and without the region of interest (Fig 5 f & g). The GEA method also identified this divergent region at the end of chromosome 2 (pos: 333357270 - 355453350) suggesting a link to ecological conditions (Fig S7). This time, this region was found to be larger than the one detected by GS (ca. 22 Mb vs 7Mb).

**Figure 5.**
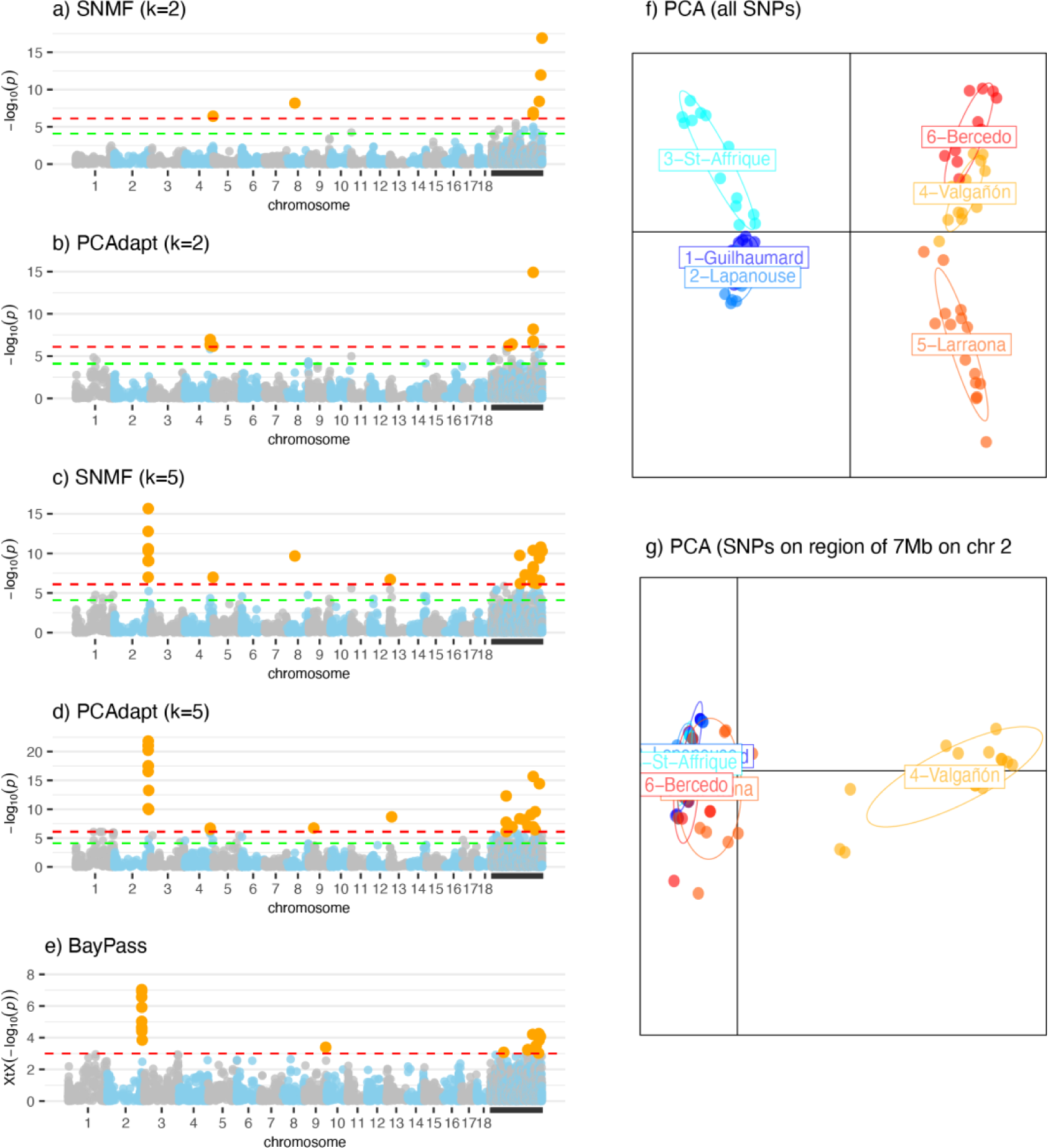
Plots (**a:e**) Manhattan plots depicting outliers SNPs detected by genome scan approaches for the *Ophrys aveyronensis* complex. Plots (**a**) and (**b**) represent outliers putatively linked to subspecies divergence, found via SNMF and PCAdapt respectively (with K=2). Plots (**c**), and (**d**) represent outliers putatively linked to populations and subspecies divergence found via SNMF, PCAdapt (with K=5) and (**e**) BayPass approaches respectively. Significant outlier loci are represented by orange dots. Labels refer to the chromosome position of outliers SNP. Red line is the genomic threshold (e.g. -log10(0.01/number of SNPs) and green line the suggestive threshold (e.g. -log10(1/number of SNPs) above which a SNP is significant. Unanchored SNPs are represented in the region beyond chromosome 19. Plots (**f,g**) Principal Component Analysis (PCA) displaying the two-firsts axes (PC1 and PC2) for (**f**) all chromosomes (86 individuals, 12744 SNPs), (**g**) region of 7Mb on chromosome 2 (86 individuals, 24 SNPs). Colors depict sampling localities. Numbers on PC1 and PC2 represent the percentage explained by the axis of the total genetic variance.

### Multiple outliers for phenotypic traits identified by reverse genetic approach

A number of SNP outliers were found to be associated with phenotypic variation, and for some these results were supported by multiple GWAS models (Table S3). Five SNPs outliers were associated with variation in “Dark green” (bin 3) and “Lichen” (bin 7) colors (Table S3)- the two traits identified as potentially under divergent selection by *P*ST-*F*ST approaches. In contrast, among the traits identified as under potentially convergent selection, only variation in the occurrence of the fatty alcohol heptacosan-1-ol (comp.21) was associated with two SNPs (Table S3). Most of there SNPs were located under unanchored regions, meaning hard to resolve and assemble regions that are often complex.

Several SNPs outliers located in the divergent region on chromosome 2 were associated with phenotypic variations (Fig 6); in particular variation in plant height, distance to first flower, occurrence of nonacosane (alkane, comp. 22) and variation in odor compounds involved in cluster n°1 (PC1, Fig 3 d). Interestingly, variation in the first dimension of odor cluster n°1 corresponds to variation in alkanes production (C21-C29) vs. other compounds such as squalene, alkylamine, ester … etc. (Fig 6, Fig S8 a and b).

**Figure 6.**
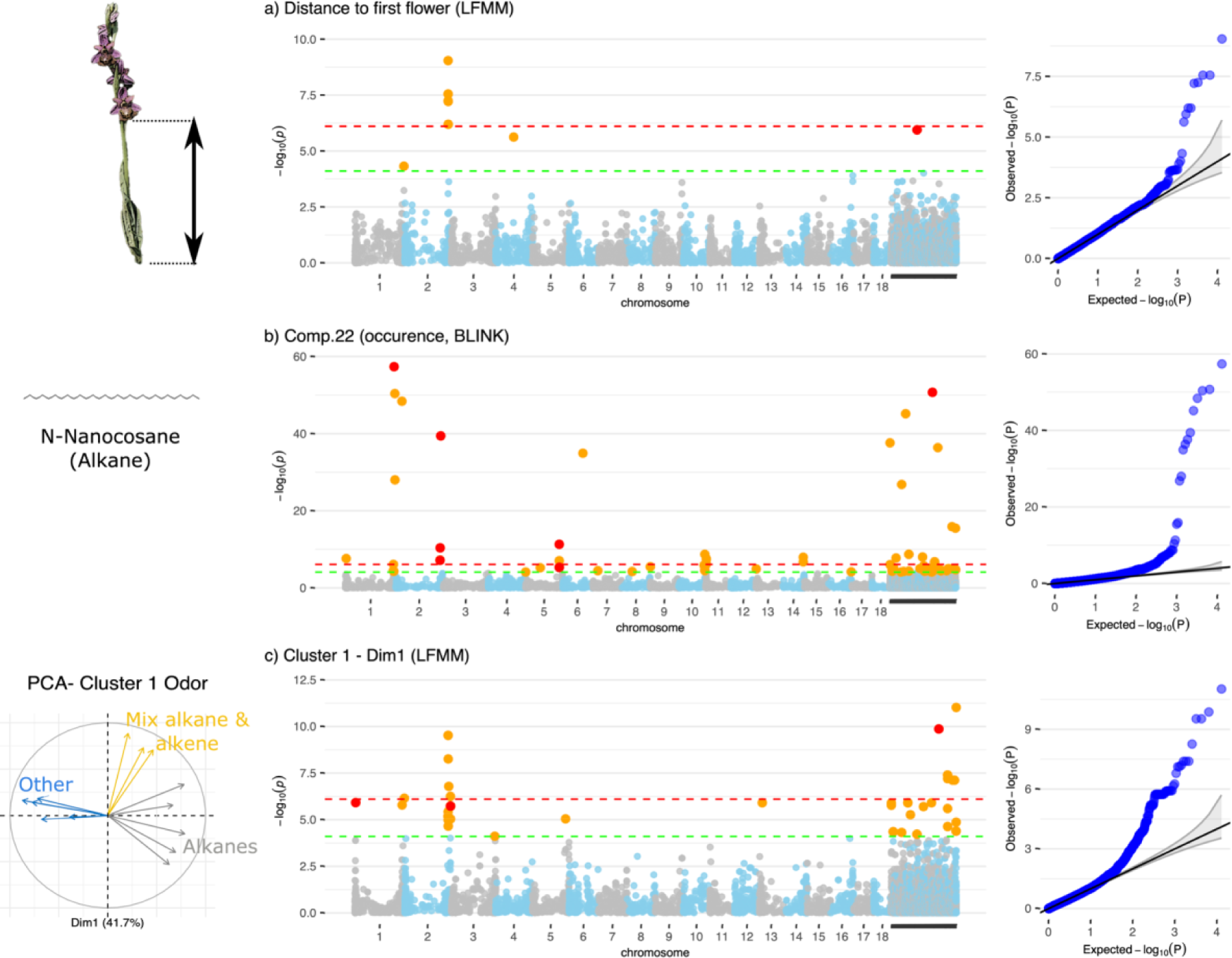
Manhattan plots depicting outliers SNPs detected by genotype-phenotype analysis (GWAS) for the *Ophrys aveyronensis* complex. Significant outlier loci are represented by orange dots, significant outlier loci found in at least two GWAS models (lfmm2, Blink and/or FarmCPU) are represented by red dots. Labels refer to the chromosome position of outliers SNP. Red line is the genomic threshold (e.g. - log10(0.01/number of SNPs) and green line the suggestive threshold (e.g. - log10(1/number of SNPs) above which a SNP is significant. Unanchored SNPs are represented in the region beyond chromosome 18.

### Candidate SNPs located in coding sequence for proteins involved in carotenoid, anthocyanin and fatty acid biosynthesis pathway

We identified 128 SNPs outliers associated with phenotypic variation that were located within or close to an annotated CDS (in a 10-kb flanking region, Table S4). Several of these CDS were known to be potentially involved in anthocyanin biosynthesis, steroid and carotenoid biosynthesis, fatty acid biosynthesis, vacuolar transporting and regulation of circadian rythm (Table 2, Fig 7). Therefore, a SNP associated with variation in ’Pink’ color (bin 6) was found within the CDS of a Flavonol synthase/flavanone 3-hydroxylase (Chr 17, evalue: 1.e-33, bitscore: 120, Table 2), an enzyme for which activity is necessary for the production of both flavonols and anthocyanins (F3H, Fig 7). Among the SNPs outliers associated to variation in odor compounds (cluster 1, alkanes comp.22, comp.7), three were located close to a CDS on chr1 encoding for a squalene/phytoene synthase. A family of enzyme known to be involved in the steroid and carotenoid pathway at the origin of carotenoid or terpenoids volatile compounds such as squalene (Fig 7). Finally, among the SNPs found in the divergent region at the end of chromosome two, one was located in the coding region of a VPS45 (Vacuolar protein sorting-associated protein 45, e-value= 1.e-31, bitscore: 115) gene.

**Figure 7:**
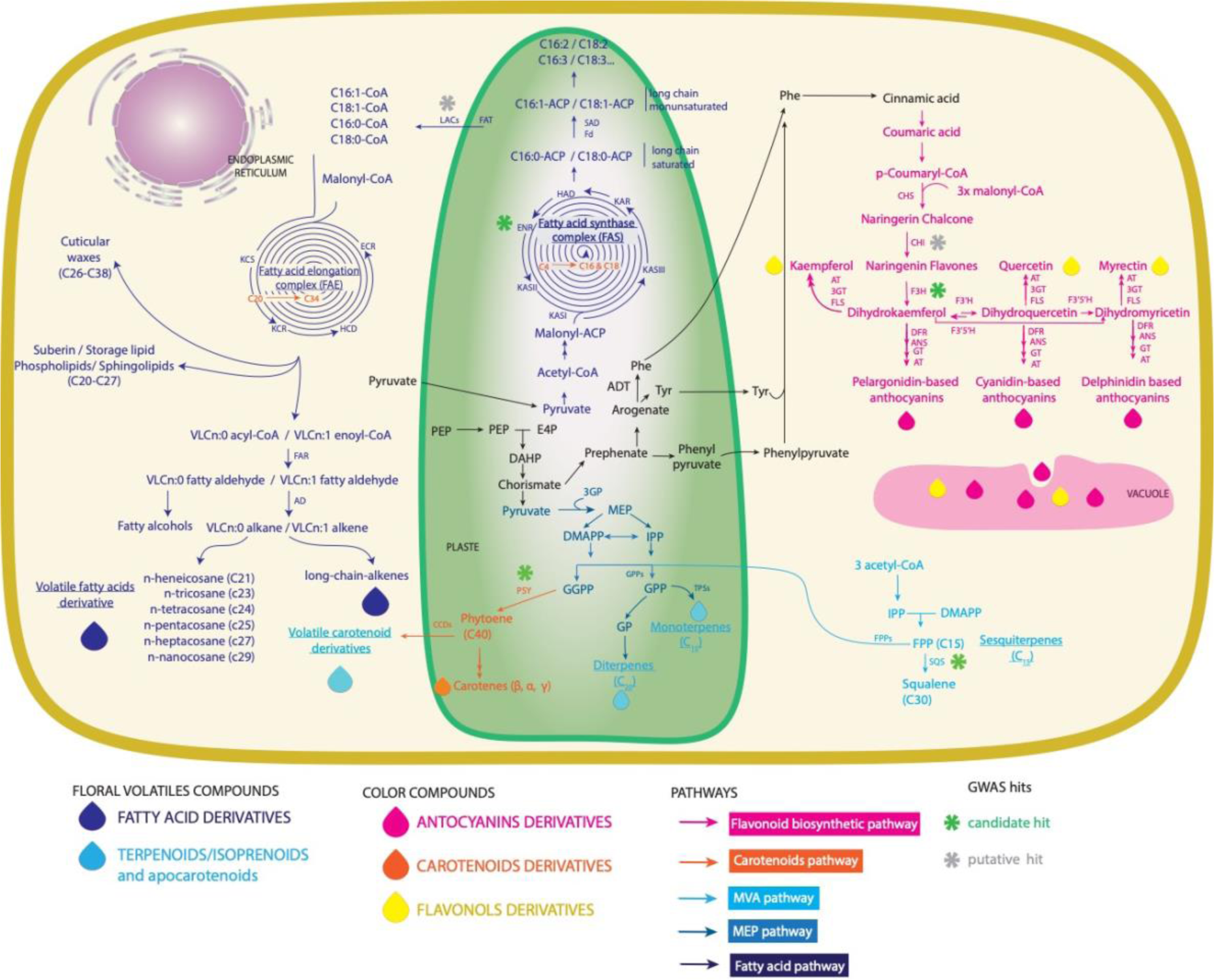
Biosynthetic pathways for pigments and volatile organic compounds (VOCs) production and storage in flowers. Adapted from Broghi *et al*. (2017). MVA: the mevalonate pathway, MEP: the methylerythritol phosphate pathway. Stars indicate enzymes for which a SNP has been found using the GWAS approach. The star is green if the enzyme is known to be involved in the trait pathway, and grey if it is not. Abbreviations : AD: aldehyde decarboxylase; ANS: anthocyanidin synthase; AT: acyltransferase CCDs: carotenoid cleavage dioxygenase; CHS: chalcone synthase; CH1: chalcone isomerase; DAHP, 3-deoxy-D-arabinoheptulosonate 7-phosphate; DMAPP, dimethylallyl pyrophosphate; DFR: dihydroflavonol 4-reductase; ENR: enoyl-ACP reductase; ECR: enoyl-CoA reductase; E4P, erythrose 4-phosphate; FAR: fatty acyl-CoA reductase; FAT: acyl-ACP thioesterase; Fd: ferredoxin; F3H: flavanone 3 hydroxylase; F3’H: flavonoid 3’hydroxylase, F3’5’H: flavonoids 3’5’- hydroxylase; FLS: flavonol synthase; FPP: farnesyl diphosphate; GGPP: geranylgeranyl diphosphate; GT: glucosyltransferase; GPP, geranyl pyrophosphate; HAD: ß-hydroacyl-ACP dehydratase; IPP, isopentenyl pyrophosphate; KAR: ß- Ketoacyl-ACP reductase; KAS (I, II, III): ß-Ketoacyl-ACP synthase (I, II, III); KCR: ß- Ketoacyl-CoA reductase; KCS: ß-Ketoacyl-CoA synthase; LACS: long-chain acyl- CoA syntgase; MEP: methylerythritol phosphate; PEP, phosphoenolpyruvate. PSY: Phytoene synthase; SAD: stearoyl/acyl-ACP desaturase; SQS: squalene synthase.

**Table 2.**
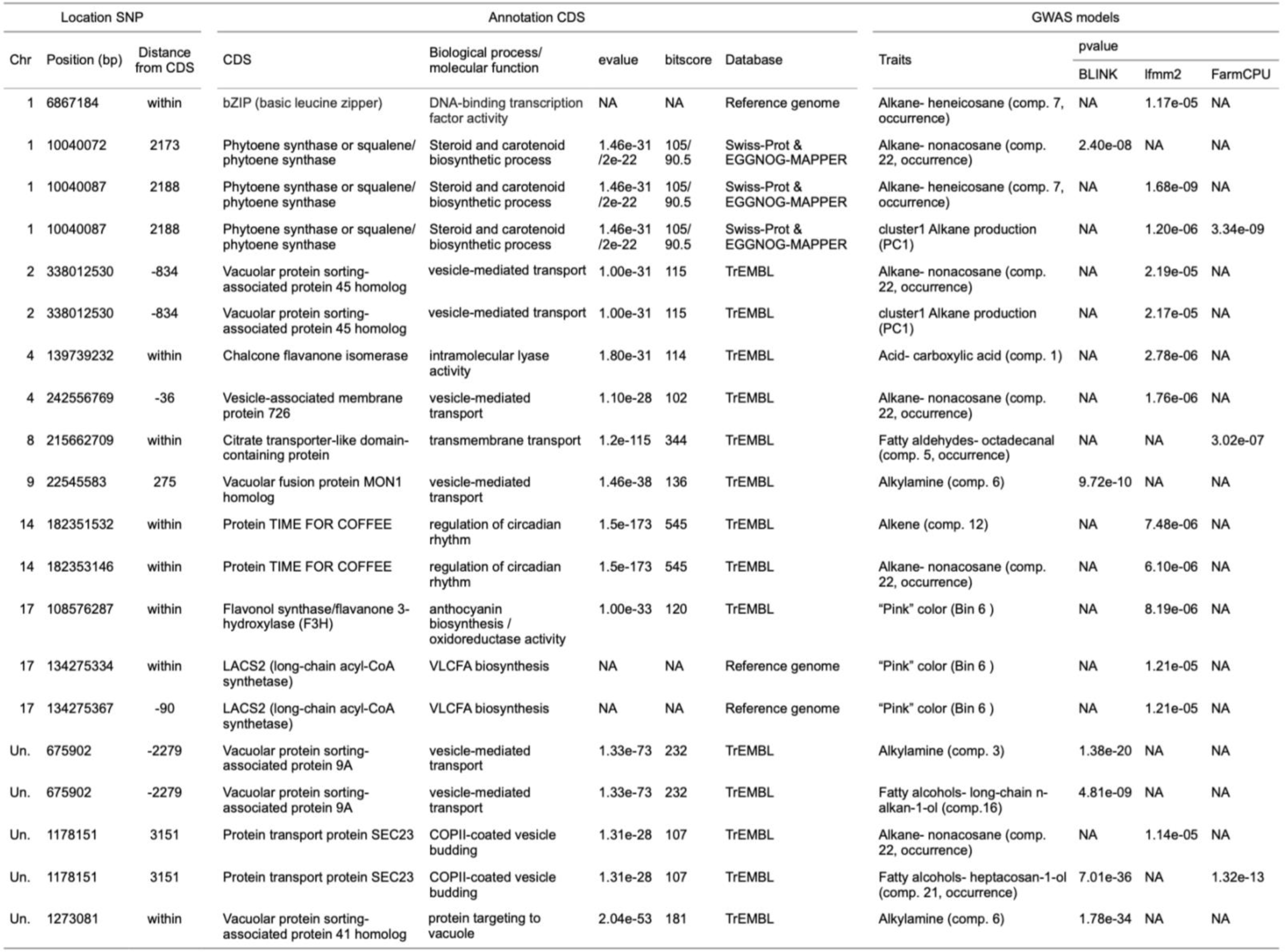
CDS of interests for SNPs identified by GWAS methods on morphological, color and odor traits in *Ophrys aveyronensis* (see complete table in appendix Table S4).

In some cases, there is a wrong match between the function of the CDS and the trait investigated in the GWAS (Fig 7). For example, two SNPs associated by GWAS with variation in ’Pink’ color (bin 6) were close (90-123bp) to the coding region of a LACS2 (Long-chain Acyl-CoA Synthase, Table 2). This enzyme is known to be involved in the fatty acid biosynthetic pathway, which produces volatile organic compounds, but is not known to be involved in anthocyanin pathway (Fig 7). At first glance, these results suggest false-positive hits in the genome-wide analysis. As we are using RAD- seq data, however, there is another possible explanation: linkage disequilibrium between multiple variants leading to the identification of a wrong target gene in the area surrounding our gene of interest. In this case, these results suggest a recurrent close proximity between color and odor genes across the genome.

In the divergent region of 7Mb at the end of chromosome 2, we identified 55 functionally annotated CDS. The GO enrichment analysis results in 18 enriched GOs (p-value < 0.01, Table S5). Among them, we observed terms associated with carbohydrate transmembrane transport (GO:0034219), acetyl-CoA biosynthetic process from pyruvate (GO:0006086), the vegetative phase change (GO:0010050), and the floral meristem determinacy (GO:0010582). In this region, a coding sequence for an Enoyl-acyl carrier protein (ACP) reductase (ENRs) was also identified (chr 2- pos: 337329689-337330517, e-value= 8.38e-103, score= 307, Table S6). This protein is known to catalyze the last step of the elongation cycle in the synthesis of fatty acids (Massengo-Tiassé & Cronan, 2009; Fig 7). When we enlarged our analyze to the region of 22 Mb identified by GEA, we found CDS associated to an AGAMOUS like MADS box protein (AGL80, chr 2 pos: 350853719-350853937, e-value = 1.57e- 28, bitscore= 109) and a Chalcone and stilbene synthase (chr2: 334453975- 334454252, e-value= 1.04e-13, bitscore= 71, Table S7).

## Discussion

Our study highlighted both a large region of genomic divergence and adaptive phenotypic traits that can potentially cause reproductive isolation.

### A genomic region of divergence on chromosome 2 associated to variation in alkane production

A large genomic region (around 7-22Mb) on chromosome 2 is strongly differentiated in *O. aveyronensis* populations; one population (in Spain) being the main cause of this divergence. A similar genomic region of divergence has recently been identified between closely-relative species of the *O. sphegodes* group in Italy (*O. exaltata*, *O. garganica* and *O. incubacea*, 20-31 Mb on Chr2, Russo *et al*. 2023). Whether this pattern is due to strong selection and/or structural variation (inversion) and/or introgression remains to be investigated. Yet, our results reinforce the idea that this region is a good candidate region for pollinator-mediated evolution in the genus *Ophrys*, and those even at the earliest stages of speciation.

Our study provides the first evidence that this genomic region was associated with phenotypic variations. In particular we detected an association with a trait cluster reflecting the production of medium and long chain n-alkanes in the floral bouquet (n- C21-n-C29). Although alkanes are common components of plants - in particular in the wax layer (Lewandowska *et al*., 2020), all of the alkanes identified here (heneicosane, tetracosane, tricosane, pentacosane, heptacosane, nonacosane) are known to be part of the sex pheromones of female insects (e.g. the moth *Orgyia leucostigma*, Grant *et al*., 1987; or the solitary bee *Andrena nigroaenea*, Schiestl *et al*., 2000). Interestingly, in the region of 7Mb, an enoyl-acyl carrier protein (ACP) reductase (ENR) was found. This gene, not annotated on the reference genome, was characterised with an annotation tool not used by Russo *et al*., (2023). ENRs catalyze the last step of the elongation cycle in the synthesis of fatty acids in the plast; extending fatty acyl chains by two carbon atoms per cycle (Massengo-Tiassé & Cronan, 2009). These results are promising, and call for further studies, especially gene expression studies, to functionally link this gene to this phenotype.

Our results showed that on this large region of divergence (7-22Mb) several genes putatively involved in the complex floral phenotype in *Ophrys* were found, suggesting the presence of supergenes (i.e. tight genetic linkage between multiple functional loci, Thompson & Jiggins, 2014; Berdan *et al*., 2022). Indeed, in the 22Mb region, besides the ENRs putatively involved in odor production, genes putatively involved in floral organ development and flavonoid biosynthesis were also detected (an AGAMOUS-like and a chalcone synthase, respectively, Li *et al*., 2020, 2022; Yeon & Kim, 2021). We did not detect any SNPs on these genes, but we must remember that the RAD- seq approach used here, only allowed us to actually sequence 0.025% of the genome (extended to 1.2% by taking into account linkage disequilibrium) (see Lowry *et al*., 2017 for a criticism concerning the use of RADseq approaches). In addition, the reference genome had ca. 1500 unanchored scaffolds (2.1% of the total assembly length, Russo *et al*., 2023), some of which were associated with phenotypic variation with SNP markers in candidate genes. Therefore, we call for further investigation of this genomic region, using long read sequencing to tackle the complexity of *Ophrys* genome. Based on all these results, this region is a strong candidate for “speciation genes” in the sexually deceptive *Ophrys*, even at an early stage of speciation.

### Adaptive differentiation in the colors that make up the macular pattern

Our study is also among the first to support an adaptive role for color in a sexually deceptive *Ophrys*. *Ophrys aveyronensis* is a species with pink/purple perianths and a greenish ‘H’ pattern on a dark brown labellum, with some variation in perianth color and blurring of the pattern between individuals (Delforge, 2016). Here we quantify this color variation and explicitly test for selection using *P*ST. Although inferences from *P*ST can be vulnerable to underlying assumptions about trait heritability (Brommer, 2011), our results are robust to a range of heritability scenarios. A signature of potential divergent selection across populations was detected on color bin 3 (dark green) and bin 7 (lichen); two related colors that form the same cluster in the trait network, and which trace the pattern on the macula. No evidence for divergent selection was found for any of the other characters. Therefore, the divergent selection could also act very early on color, such as what is expected for olfactory compounds (Baguette *et al*., 2020). Unfortunately, it was not possible to test for divergence/convergence of the most important olfactory compounds in particular alkanes and alkenes (because of lack of model robustness).

Our results relaunch the debate on the role of the pattern on the macula. Indeed, visual cues are classically considered of secondary importance for pollinator attraction in the sexually deceptive *Ophrys* compared to olfactory cues (Schiestl *et al*., 2000; Schiestl, 2005; Stökl *et al*., 2008; Vereecken & Schiestl, 2009). Several authors suggested that visual cues would be subject to less evolutionary potential in pollinator shifts (Baguette *et al*., 2020). In the literature, the patterns on the labellum were thought to play a role in pollinator attraction by mimicking the wings or body markings of the female pollinator, but this expectation was not supported by field observations. In *Ophrys heldreichii* - a species with a complex and variable macula - no evidence was found that its pollinators (*Eucera berlandi*) discriminate between flowers with and without a pattern on the labellum (Streinzer *et al*., 2010). Instead, it has been hypothesized that dissatisfied pollinators would learn the pattern to avoid re-visits (Stejskal *et al*., 2015); *Ophrys* would “use” the labellum pattern as a strategy to increase outcrossing. How this selective force would drive divergence between populations is intriguing and requires further investigation, including behavioral experiments.

### Convergence (or canalization) on morphological trait and on fatty alcohols odor compounds

Our study revealed some trait conservatism among populations of *Ophrys aveyronensis*, meaning little or no variation between populations. Indeed, flower morphological traits showed low variability compared to other traits and formed independent nodes, suggesting stability and conservatism. This result is consistent with Berg’s hypothesis, which states that flower size in insect-pollinated plants should be selected to remain constant regardless of the size of the vegetative structure, so that flowers match pollinator size for efficient pollination (Berg, 1960). This result is reassured by the evidence of significantly less phenotypic variation in the labellum width and petal width traits than would be expected by genetic drift, with a significant *P*ST < *F*ST pattern between populations. The classical hypothesis for *P*ST < *F*ST is that selection should have approximately the same selective optimum in all populations (Lamy *et al*., 2012). While it is difficult to imagine a mechanism selecting for a given petal width, selection for a given labellum size or shape has already been hypothesized in the literature. The labellum shape and size were expected to morphologically mimic the females of the pollinator species (Peakall, 1990), or to be mechanically conform to the morphology of the male pollinator to improve copulation and/or contact with pollinia (de Jager & Peakall, 2016) with flowers form components used as gripping points to be more effective in ensuring pollinia transfer (Rakosy *et al*., 2017). Yet, an alternative hypothesis to trait conservatism is canalisation, where the genetic backgrounds of populations share the same genetic constraint (low or no genetic variance) rather than the same optima (Hansen & Houle, 2004; Lamy *et al*., 2012). While the convergent selection hypothesis for labellum size is appealing, it is not possible to discriminate between the two hypotheses here. Elucidating the genomic architecture of these traits remains a major challenge in *Ophrys*, despite important progress in orchids (Li *et al*., 2022).

Interestingly, the same pattern of trait conservatism was observed for three odor traits, all fatty alcohols. While odor traits as a whole showed high variability and formed interconnected clusters of covariant traits, these three fatty alcohols displayed a different pattern. They showed low variance, formed independent clusters and exhibited a pattern of *P*ST < *F*ST between populations. To our knowledge, the role of fatty alcohols in pollinator attraction is relatively less understood than a mixture of alkenes and alkanes (Peakall, 2023). Our result suggests a potential role in pollinator attraction or a genetic limitation that remains to be investigated.

### Standing variation in color and odor not linked to signature of selection

Our study suggests standing genetic variation in candidate genes associated with color and odor traits involved in the complex floral phenotype of *Ophrys*. Variation in purple color on petal and sepal (bin6- pink) was associated with a SNPs marker for a flavone synthase, a protein known to be involved in anthocyanin biosynthesis in the flower (Orteu & Jiggins, 2020; Yeon & Kim, 2021). A variation in a long chain n-alkene (compound 12) was associated with a marker on an acetyl-CoA-C acetyltransferase, an enzyme putatively involved in the fatty acid biosynthetic pathway and considered a good candidate for odor production in *Ophrys* (Russo *et al*., 2023). Our study suggested a pleiotropy between color and odor, found through the trait network and supported by GWAS analysis, with odor traits linked to genes putatively involved in color production and vice versa (Yeon & Kim, 2021). While none of these traits has yet shown a signature of adaptive differentiation between populations (i.e., no evidence for a signature for bin 6 and no model convergence for alkene comp. 12), adaptation via standing genetic variation is thought to be particularly important in cases of rapid adaptation (Barrett & Schlüter, 2008). Our study demonstrated the interest of integrative approaches (e.g., reverse and forward genomic approaches, completed with *P*ST-*F*ST comparison) to distinguish between standing genetic variation for a phenotype and a target of selection (Barrett & Hoekstra, 2011; Bomblies & Peichel, 2022; Joffard *et al*., 2022).

## Conclusion

Our study reveals patterns that often become invisible over time, i.e., the geographic mosaic of traits under selection and the early appearance of strong genomic divergence driven by a few populations. Contrary to our initial expectation, a signature of divergent selection across populations was found for color, but no evidence was found for olfactory compounds (Baguette *et al*., 2020). As expected following Berg’s theory (1960), floral morphological traits showed low variability and independence, indicating stability and conservatism. Odor and color traits, on the other hand, showed high variability and formed interconnected clusters, suggesting rapid adaptive potential. Finally, we found evidence for variation in odors but not associated with SAD types genes (Schlüter *et al*., 2011; Xu *et al*., 2012, 2017). Instead, our candidate genes were involved further up in the fatty acid or in the steroid biosynthetic pathways.

## Acknowledgments

We thank Jean-Luc Roux and Daniel Vizcaïno for the information provided about sampling sites in Spain.

## Author Contributions

Conceptualisation, J.A.M.B., A.G. B.S. ; data acquisition, A.G., R.B., B.S. and J.A.M.B.; methodology, J.A.M.B., B.S., A.G., D.N. and M.B.; odor compounds identification: N.B. & D.N., formal analysis, A.G and J.A.M.B; writing—original draft preparation, A.G.; writing—review and editing, A.G., J.A.M.B., B.S., R.B., D.N. and M.B.; supervision, J.A.M.B.; project administration, J.A.M.B.; funding acquisition, J.A.M.B., B.S.; All authors have read and agreed to the published version of the manuscript.

## Data Availability Statement

All the data sets (phenotypic data, picture and metadata input files) used in the analyses presented in this article and the R code protocols used to run these analyses with these data sets are already freely accessible on the ZENODO. Sequencing data have been submitted to the European Nucleotide Archive (ENA; https://www.ebi.ac.uk/ena/browser/home) under Study with primary accession n°PRJEB61037 (and secondary accession number ERP146119) and samples accession n°ERS14864169 (SAMEA112857509) to n°ERS14864254 (SAMEA112857594).

## References

1. Alexa A, Rahnenfuhrer J. 2023. topGO: Enrichment Analysis for Gene Ontology.

2. Ayasse M, Schiestl FP, Paulus HF, Fstedt CL, Hansson B, Ibarra F, Francke W. 2000. Evolution of reproductive strategies in the sexually deceptive Orchid *Ophrys sphegodes*: How does flower-specific variation of odor signals influence reproductive success? Evolution: 12.

3. Ayasse M, Schiestl FP, Paulus HF, Ibarra F, Francke W. 2003. Pollinator attraction in a sexually deceptive orchid by means of unconventional chemicals. Proceedings of the Royal Society of London. Series B: Biological Sciences 270: 517–522.

4. Baguette M, Bertrand JAM, Stevens VM, Schatz B. 2020. Why are there so many bee-orchid species? Adaptive radiation by intra-specific competition for mnesic pollinators. Biological Reviews 95: 1630–1663.

5. Barrett RDH, Hoekstra HE. 2011. Molecular spandrels: tests of adaptation at the genetic level. Nature Reviews Genetics 12: 767–780.

6. Barrett RDH, Schlüter D. 2008. Adaptation from standing genetic variation. Trends in Ecology & Evolution 23: 38–44.

7. Barton K. 2009. Mu-MIn: Multi-model inference. R Package.

8. Bates D, Mächler M, Bolker B, Walker S. 2015. Fitting linear mixed-effects models using lme4. Journal of Statistical Software 67.

9. Benito Ayuso J. 2019. Estudios sobre polinizacion en el genero Ophrys (Orchidaceae). Flora Montiberica 74: 32–37.

10. Benjamini Y, Hochberg Y. 1995. Controlling the False Discovery Rate: A Practical and Powerful Approach to Multiple Testing. Journal of the Royal Statistical Society: Series B (Methodological) 57: 289–300.

11. Berdan EL, Flatt T, Kozak GM, Lotterhos KE, Wielstra B. 2022. Genomic architecture of supergenes: connecting form and function. Philosophical Transactions of the Royal Society B: Biological Sciences 377: 20210192.

12. Berg RL. 1960. The Ecological Significance of Correlation Pleiades. Evolution 14: 171--180.

13. Bertrand JAM, Delahaie B, Bourgeois YXC, Duval T, García-Jiménez R, Cornuault J, Pujol B, Thébaud C, Milá B. 2016. The role of selection and historical factors in driving population differentiation along an elevational gradient in an island bird. Journal of Evolutionary Biology 29: 824–836.

14. Bomblies K, Peichel CL. 2022. Genetics of adaptation. Proceedings of the National Academy of Sciences 119: e2122152119.

15. Breitkopf H, Onstein RE, Cafasso D, Schlüter PM, Cozzolino S. 2015. Multiple shifts to different pollinators fuelled rapid diversification in sexually deceptive Ophrys orchids. New Phytologist 207: 377–389.

16. Brommer JE. 2011. Whither Pst? The approximation of Qst by Pst in evolutionary and conservation biology: Whither Pst? Journal of Evolutionary Biology 24: 1160– 1168.

17. Broghi M, Fernie A, Schiestl F, Bowmeester H. 2017 The Sexual Advantage of Looking, Smelling, and Tasting Good: The Metabolic Network that Produces Signals for Pollinators. Trends in Plant Science 22: 338–350.

18. Cantalapiedra CP, Hernández-Plaza A, Letunic I, Bork P, Huerta-Cepas J. 2021. eggNOG-mapper v2: Functional Annotation, Orthology Assignments, and Domain Prediction at the Metagenomic Scale. Molecular Biology and Evolution 38: 5825– 5829.

19. Catchen JM, Amores A, Hohenlohe PA, Fresko W, Postlethwait JH. 2011. Stacks: building and genotyping loci de novo from short-read sequences, G3 Genes|Genomes|Genetics, Volume 1, Issue 3, 1 August 2011, Pages 171–182, 10.1534/g3.111.000240

20. Catchen JM, Hohenlohe PA, Bassham S, Amores A, Cresko WA. 2013. Stacks: an analysis tool set for population genomics. Molecular Ecology 22: 3124–3140. doi: 10.1111/mec.12354

21. Csárdi G, Nepusz T, Müller K, Horvát S, Traag V, Zanini F, Noom D. 2023. igraph for R: R interface of the igraph library for graph theory and network analysis.

22. Delforge P. 2016. Orchidées d’Europe, d’Afrique du Nord et du Proche-Orient: la bible des orchidophiles, plus de 600 espèces et de nombreuses variétés et illustrées. Paris: Delachaux et Niestlé.

23. Frichot E, François O. 2015. LEA: An R package for landscape and ecological association studies (B O’Meara, Ed.). Methods in Ecology and Evolution 6: 925–929.

24. Gervasi DDL, Selosse M-A, Sauve M, Francke W, Vereecken NJ, Cozzolino S, Schiestl FP. 2017. Floral scent and species divergence in a pair of sexually deceptive orchids. Ecology and Evolution 7: 6023–6034.

25. Gibert A, Buscail R, Baguette M, Fraïsse C, Roux C, Schatz B, Bertrand JAM. 2023. Climate change promoted allopatric divergence and explains the current disjunct geographic distribution of the *Ophrys aveyronensis* species complex (Orchidaceae). bioRxiv: 2023.04.27.538532.

26. Gibert A, Louty F, Buscail R, Baguette M, Schatz B, Bertrand JAM. 2022. Extracting Quantitative Information from images taken in the wild: a case study of two vicariants of the *Ophrys aveyronensis* species complex. Diversity 14: 400.

27. Grant GG, Frech D, MacDonald L, Slessor KN, King GGS. 1987. Copulation releaser pheromone in body scales of female whitemarked tussock moth, *Orgyia leucostigma* (Lepidoptera: Lymantriidae): Identification and behavioral role. Journal of Chemical Ecology 13: 345–356.

28. Grueber CE, Nakagawa S, Laws RJ, Jamieson IG. 2011. Multimodel inference in ecology and evolution: challenges and solutions: Multimodel inference. Journal of Evolutionary Biology 24: 699–711.

29. Hansen TF, Houle D. 2004. Evolvability, Stabilizing Selection, and the Problem of Stasis. In: Pigliucci M, Preston K, eds. Phenotypic Integration: Studying the Ecology and Evolution of Complex Phenotypes. Oxford University Press.

30. Harrison XA, Donaldson L, Correa-Cano ME, Evans J, Fisher DN, Goodwin CED, Robinson BS, Hodgson DJ, Inger R. 2018. A brief introduction to mixed effects modelling and multi-model inference in ecology. PeerJ 6: e4794.

31. Hartig F. 2020. DHARMa: Residual Diagnostics for Hierarchical (Multi-Level / Mixed) Regression Models. R package.

32. He N, Li Y, Liu C, Xu L, Li M, Zhang J, He J, Tang Z, Han X, Ye Q, et al. 2020. Plant trait networks: improved resolution of the dimensionality of adaptation. Trends in Ecology & Evolution 35: 908–918.

33. Hermosilla CE, Sabando J. 1998. Notas sobre orquídeas (V). Estudios del Museo de Ciencias Naturales de Álava 13: 123–156.

34. Hermosilla CE, Soca R. 1999. Distribuzione di *Ophrys aveyronensis* (J.J. Wood) Delforge (Orchidaceae) e rassegna dei suoi ibridi. Caesiana. Caesiana 13: 31–38.

35. Hijmans RJ, Cameron SE, Parra JL, Jones PG, Jarvis A. 2005. Very high resolution interpolated climate surfaces for global land areas. International Journal of Climatology 25: 1965–1978.

36. Huang M, Liu X, Zhou Y, Summers RM, Zhang Z. 2019. BLINK: a package for the next level of genome-wide association studies with both individuals and markers in the millions. GigaScience 8: giy154.

37. de Jager ML, Peakall R. 2016. Does morphology matter? An explicit assessment of floral morphology in sexual deception. Functional Ecology 30: 537–546.

38. Joffard N, Buatois B, Schatz B. 2016. Integrative taxonomy of the fly orchid group: insights from chemical ecology. The Science of Nature 103: 77.

39. Joffard N., Buatois B., Arnal V., Véla E., Montgelard C., Schatz B. 2022. Delimiting species in the taxonomically challenging orchid section Pseudophrys: Bayesian analyses of genetic and phenotypic data. Front. Ecol. Evol. 10.3389/fevo.2022.1058550

40. Jombart T, Ahmed I. 2011. adegenet 1.3-1 : new tools for the analysis of genome- wide SNP data. Bioinformatics 27: 3070–3071.

41. Lai J, Zou Y, Zhang J, Peres-Neto PR. 2022. Generalizing hierarchical and variation partitioning in multiple regression and canonical analyses using the rdacca.hp R package. Methods in Ecology and Evolution 13: 782–788.

42. Lamy J-B, Plomion C, Kremer A, Delzon S. 2012. QST < FST as a signature of canalization. Molecular Ecology 21: 5646–5655.

43. Lê Cao K-A, Boitard S, Besse P. 2011. Sparse PLS discriminant analysis: biologically relevant feature selection and graphical displays for multiclass problems. BMC Bioinformatics 12: 253.

44. Leinonen T, O’Hara RB, Cano JM, Merilä J. 2008. Comparative studies of quantitative trait and neutral marker divergence: a meta-analysis: QST - FST meta-analysis. Journal of Evolutionary Biology 21: 1–17.

45. Lewandowska M, Keyl A, Feussner I. 2020. Wax biosynthesis in response to danger: its regulation upon abiotic and biotic stress. New Phytologist 227: 698–713.

46. Li Y, Zhang B, Yu H. 2022. Molecular genetic insights into orchid reproductive development (R Melzer, Ed.). Journal of Experimental Botany 73: 1841–1852.

47. Li B-J, Zheng B-Q, Wang J-Y, Tsai W-C, Lu H-C, Zou L-H, Wan X, Zhang D-Y, Qiao H-J, Liu Z-J, et al. 2020. New insight into the molecular mechanism of colour differentiation among floral segments in orchids. Communications Biology 3: 89.

48. Liaw, Wiener. 2015. Package ‘randomForest’: Breiman and Cutler’s random forests for classification and regression.

49. Liu X, Huang M, Fan B, Buckler ES, Zhang Z. 2016. Iterative usage of fixed and random effect models for powerful and efficient genome-wide association studies. PLOS Genetics 12: e1005767.

50. Lowry DB, Hoban S, Kelley JL, Lotterhos KE, Reed LK, Antolin MF, Storfer A. 2017. Breaking RAD: an evaluation of the utility of restriction site-associated DNA sequencing for genome scans of adaptation. Molecular Ecology Resources 17: 142– 152.

51. Mant J, Peakall R, Schiestl FP. 2005. Does selection on floral odor promote differentiation among populations and species of the sexually deceptive orchid genus ophrys? Evolution 59: 1449.

52. Massengo-Tiassé RP, Cronan JE. 2009. Diversity in enoyl-acyl carrier protein reductases. Cellular and molecular life sciences: CMLS 66: 1507–1517.

53. Newman MEJ. 2006. Modularity and community structure in networks. Proceedings of the National Academy of Sciences 103: 8577–8582.

54. Newman MEJ, Girvan M. 2004. Finding and evaluating community structure in networks. Physical Review E 69: 026113.

55. Oksanen J, Kindt R, Legendre P, O’Hara B, Simpson G, Solymos P, Stevens M, Wagner H. 2009. The VEGAN Package: community ecology package.

56. Orteu A, Jiggins CD. 2020. The genomics of coloration provides insights into adaptive evolution. Nature Reviews Genetics 21: 461–475.

57. Paulus HF. 2017. Zur bestäubungsbiologie der gattung Ophrys in nordspanien: freilandstudien an *Ophrys aveyronensis, O. subinsectifera, O. riojana, O. vasconica* und *O. forestieri*. J. Eur. Orch. 49: 427–471.

58. Peakall R. 1990. Responses of male *Zaspilothynnus trilobatus* turner wasps to females and the sexually deceptive orchid it pollinates. Functional Ecology 4: 159– 167.

59. Peakall R. 2023. Pollination by sexual deception. Current Biology 33: R489–R496.

60. Pélabon C, Hilde CH, Einum S, Gamelon M. 2020. On the use of the coefficient of variation to quantify and compare trait variation. Evolution Letters 4: 180–188.

61. R Core Team. 2018. R: a language and environment for statistical computing. Vienna, Austria: R Foundation for Statistical Computing.

62. Russo A, Alessandrini M, El Baidouri M, Frei D, Galise T, Gaidusch L, Oerter H, Garcia Morales S, Potente G, Tian Q, Smetanin S, Bertrand J, Onstein R, Panaud O, Frey J, Cozzolino S, Wicker T, Xu S, Grossniklaus U, Schlüter P. 2023. The genome of the early spider-orchid *Ophrys sphegodes* provides insights into sexual deception and adaptation to pollinators. Research Square. DOI: 10.21203/rs.3.rs-3463148/v1

63. Rakosy D, Cuervo M, Paulus HF, Ayasse M. 2017. Looks matter: changes in flower form affect pollination effectiveness in a sexually deceptive orchid. 30: 1978–1993.

64. Rohart F, Gautier B, Singh A, Cao K-AL. 2017. mixOmics: An R package for ‘omics feature selection and multiple data integration. PLOS Computational Biology 13: e1005752.

65. Schiestl FP. 2005. On the success of a swindle: pollination by deception in orchids. Naturwissenschaften 92: 255–264.

66. Schiestl FP, Ayasse M. 2002. Do changes in floral odor cause speciation in sexually deceptive orchids? Plant Systematics and Evolution 234: 111–119.

67. Schiestl FP, Ayasse M, Paulus HF, Löfstedt C, Hansson BS, Ibarra F, Francke W. 2000. Sex pheromone mimicry in the early spider orchid (*Ophrys sphegodes*): patterns of hydrocarbons as the key mechanism for pollination by sexual deception. Journal of Comparative Physiology A: Sensory, Neural, and Behavioral Physiology 186: 567–574.

68. Schiestl FP, Schlüter PM. 2009. Floral isolation, specialized pollination, and pollinator behavior in orchids. Annual Review of Entomology 54: 425–446.

69. Schlüter PM. 2018. The magic of flowers or: speciation genes and where to find them. American Journal of Botany 105: 1957–1961.

70. Schlüter D, Rieseberg LH. 2022. Three problems in the genetics of speciation by selection. Proceedings of the National Academy of Sciences 119: e2122153119.

71. Schlüter PM, Xu S, Gagliardini V, Whittle E, Shanklin J, Grossniklaus U, Schiestl FP. 2011. Stearoyl-acyl carrier protein desaturases are associated with floral isolation in sexually deceptive orchids. Proceedings of the National Academy of Sciences 108: 5696–5701.

72. Seeholzer GF, Brumfield RT. 2018. Isolation by distance, not incipient ecological speciation, explains genetic differentiation in an Andean songbird (Aves: Furnariidae: *Cranioleuca antisiensis*, Line-cheeked Spinetail) despite near threefold body size change across an environmental gradient. Molecular Ecology 27: 279–296.

73. Stejskal K, Streinzer M, Dyer A, Paulus HF, Spaethe J. 2015. Functional significance of labellum pattern variation in a sexually deceptive orchid (*Ophrys heldreichii*): Evidence of Individual Signature Learning Effects (RM Borges, Ed.). PLOS ONE 10: e0142971.

74. Stökl J, Schlüter PM, Stuessy TF, Paulus HF, Assum G, Ayasse M. 2008. Scent variation and hybridization cause the displacement of a sexually deceptive orchid species. American Journal of Botany 95: 472–481.

75. Streinzer M, Ellis T, Paulus HF, Spaethe J. 2010. Visual discrimination between two sexually deceptive Ophrys species by a bee pollinator. Arthropod-Plant Interactions 4: 141–148.

76. The UniProt Consortium. 2023. UniProt: the Universal Protein Knowledgebase in 2023. Nucleic Acids Research 51: D523–D531.

77. Thompson MJ, Jiggins CD. 2014. Supergenes and their role in evolution. Heredity113: 1–8.

78. Vereecken NJ, Schiestl FP. 2009. On the roles of colour and scent in a specialized floral mimicry system. Annals of Botany 104: 1077–1084.

79. Wang J, Zhang Z. 2021. GAPIT Version 3: Boosting Power and Accuracy for Genomic Association and Prediction. Genomics, Proteomics & Bioinformatics 19: 629–640.

80. Weir BS, Cockerham CC. 1984. Estimating F-Statistics for the Analysis of Population Structure. Evolution 38: 1358.

81. Weller HI, Westneat MW. 2019. Quantitative color profiling of digital images with earth mover’s distance using the R package colordistance. PeerJ. 6;7:e6398. doi: 10.7717/peerj.6398.

82. Wood JJ. 1983. Two New Combinations in Ophrys (Orchidaceae). Kew Bulletin 38: 135.

83. Xu H, Bohman B, Wong DCJ, Rodriguez-Delgado C, Scaffidi A, Flematti GR, Phillips RD, Pichersky E, Peakall R. 2017. Complex Sexual Deception in an Orchid Is Achieved by Co-opting Two Independent Biosynthetic Pathways for Pollinator Attraction. Current Biology 27: 1867–1877.e5.

84. Xu S, Schlüter PM, Grossniklaus U, Schiestl FP. 2012. The genetic basis of pollinator adaptation in a sexually deceptive orchid (Ps Schnable, Ed.). PLoS Genetics 8: e1002889.

85. Yeon JY, Kim WS. 2021. Biosynthetic Linkage between the Color and Scent of Flowers: A Review. Horticultural Science & Technology 39: 697–713.

