## Supplementary material for "Floral phenotypic divergence and genomic insights in an *Ophrys* orchid: Unraveling early speciation processes"

Article acceptance date:

The following Supporting Information is available for this article:

**Fig. S1** Boxplots of (**a:f**) plant-level morphological traits, (**g:n**) floral morphological traits, (**o:v**) color traits and (**w:at**) odor traits.

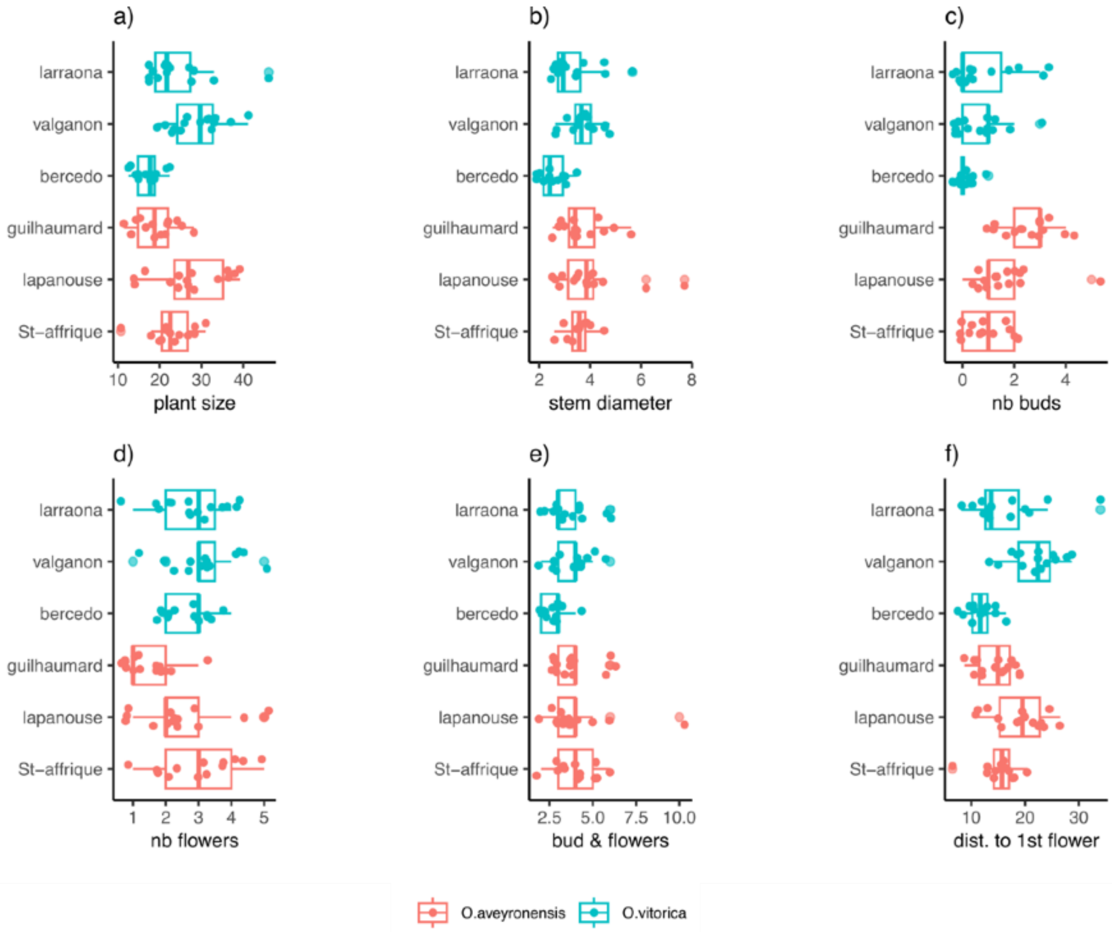

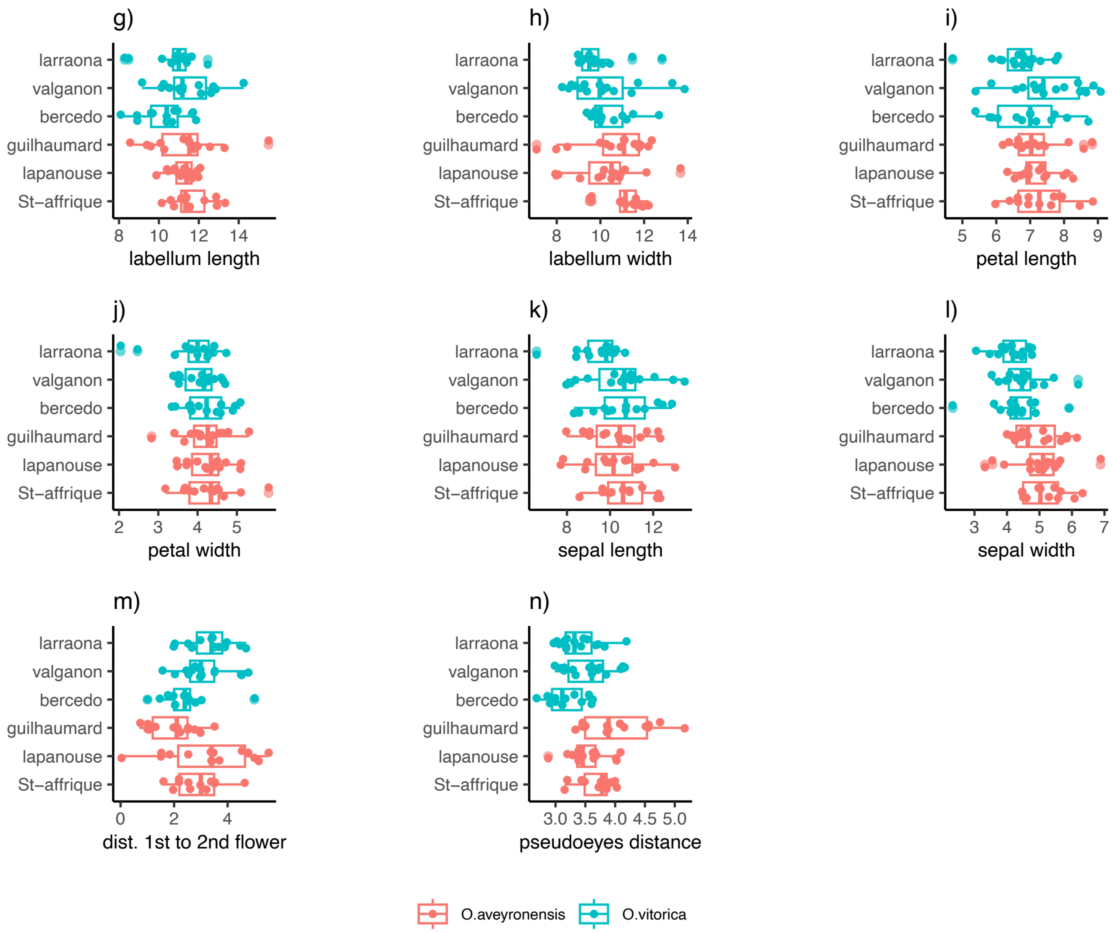
g)
h)
i)
j)
k)
l)
m)
n)
larraona
valganon
bercedo
guilhaumard
lapanouse
St–affrique
labellum length
labellum width
petal length
petal width
sepal length
sepal width
dist. 1st to 2nd flower
pseudoeyes distance
O.aveyronensis
O.vitorica

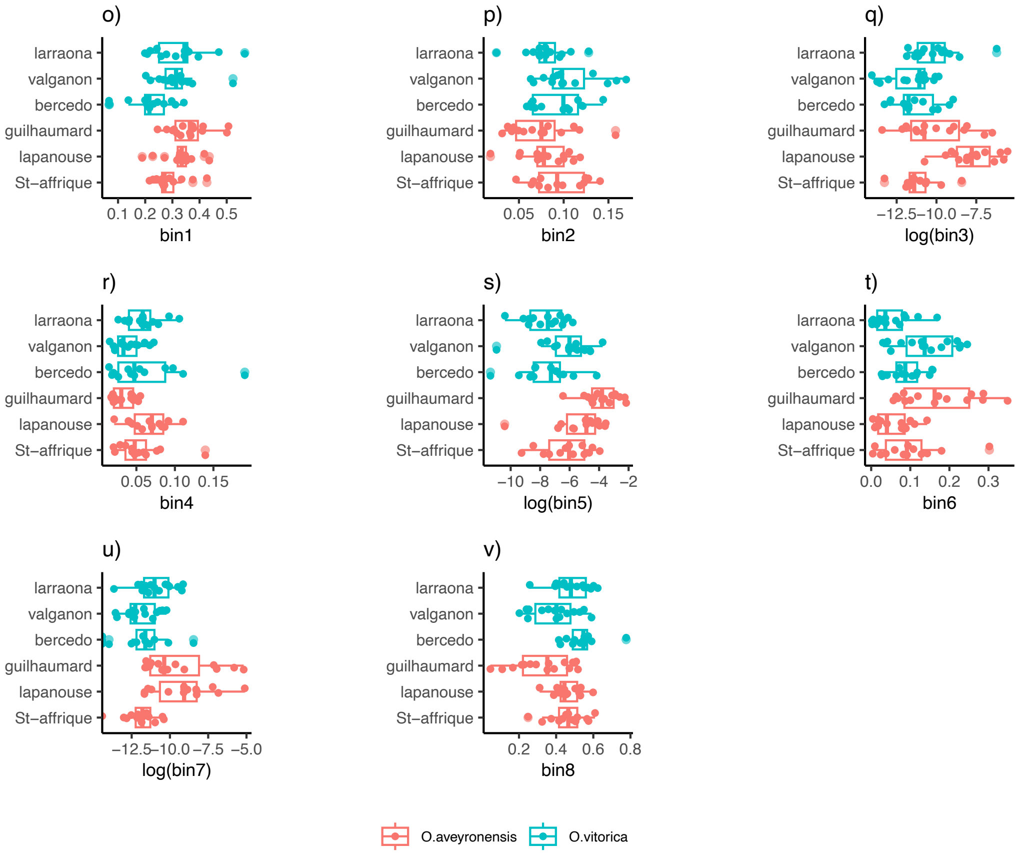
o)
p)
q)
r)
s)
t)
u)
v)
larraona
valganon
bercedo
guilhaumard
lapanouse
St–affrique
bin1
bin2
log(bin3)
bin4
log(bin5)
bin6
log(bin7)
bin8
O.aveyronensis
O.vitorica

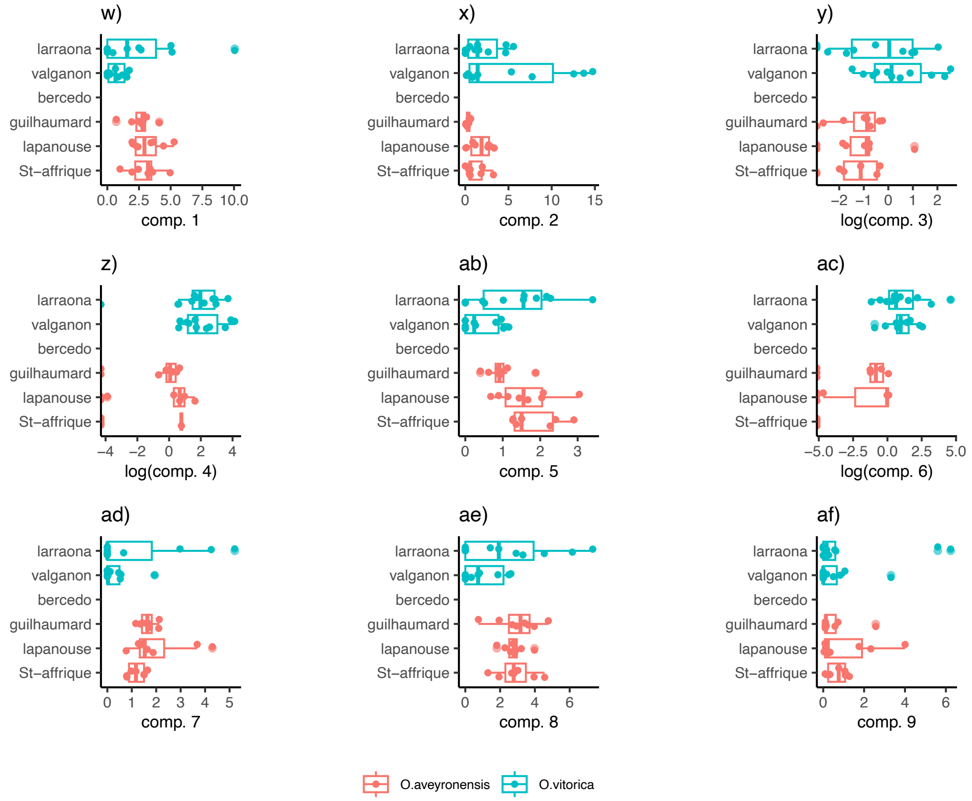
w)
x)
y)
z)
ab)
ac)
ad)
ae)
af)
larraona
valganon
bercedo
guilhaumard
lapanouse
St–affrique
comp. 1
comp. 2
log(comp. 3)
log(comp. 4)
comp. 5
log(comp. 6)
comp. 7
comp. 8
comp. 9
O.aveyronensis
O.vitorica

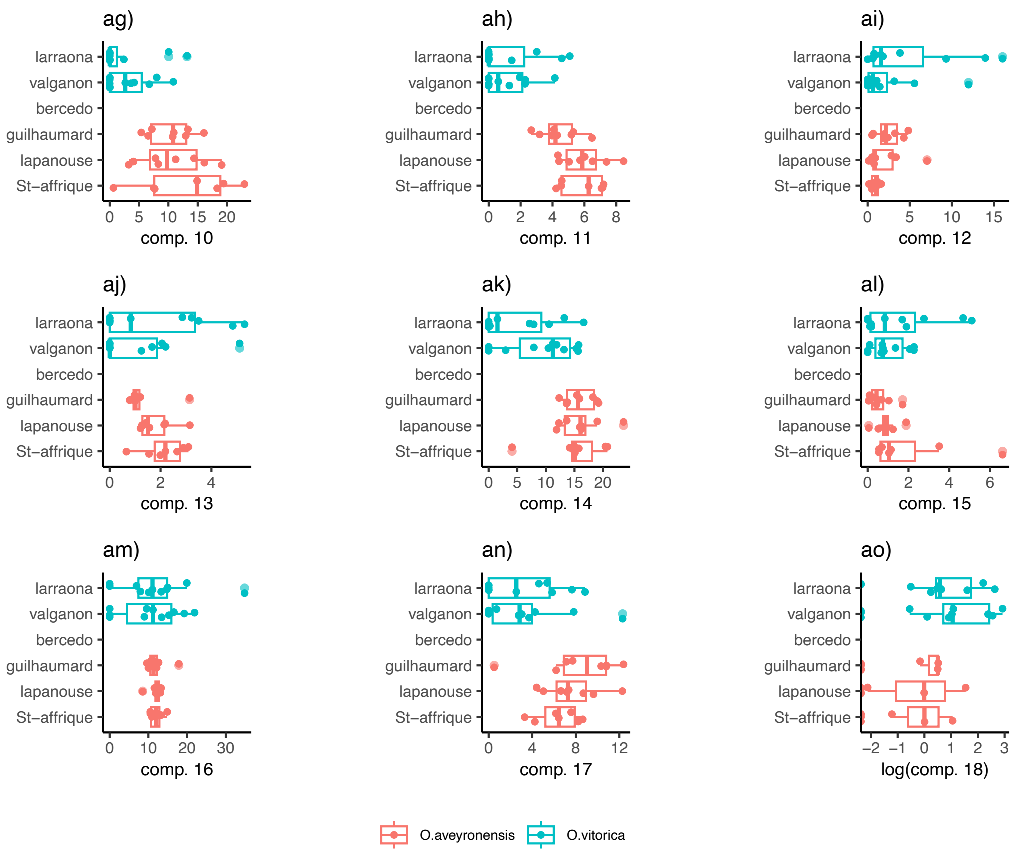
ag)
ah)
ai)
aj)
ak)
al)
am)
an)
ao)
larraona
valganon
bercedo
guilhaumard
lapanouse
St–affrique
comp. 10
comp. 11
comp. 12
comp. 13
comp. 14
comp. 15
comp. 16
comp. 17
log(comp. 18)
O.aveyronensis
O.vitorica

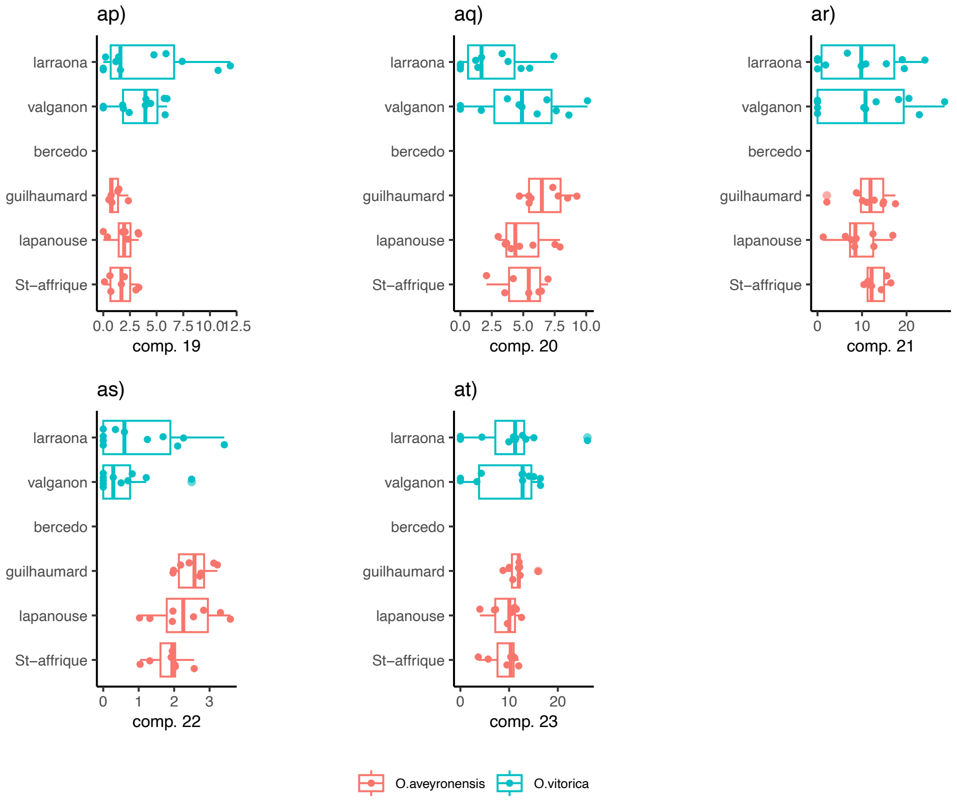
ap)
aq)
ar)
as)
at)
larraona
valganon
bercedo
guilhaumard
lapanouse
St–affrique
comp. 19
comp. 20
comp. 21
comp. 22
comp. 23
O.aveyronensis
O.vitorica

**Fig. S2** Distribution of genome-wide SNPs in *Ophrys aveyronensis* chromosome-scale genome.

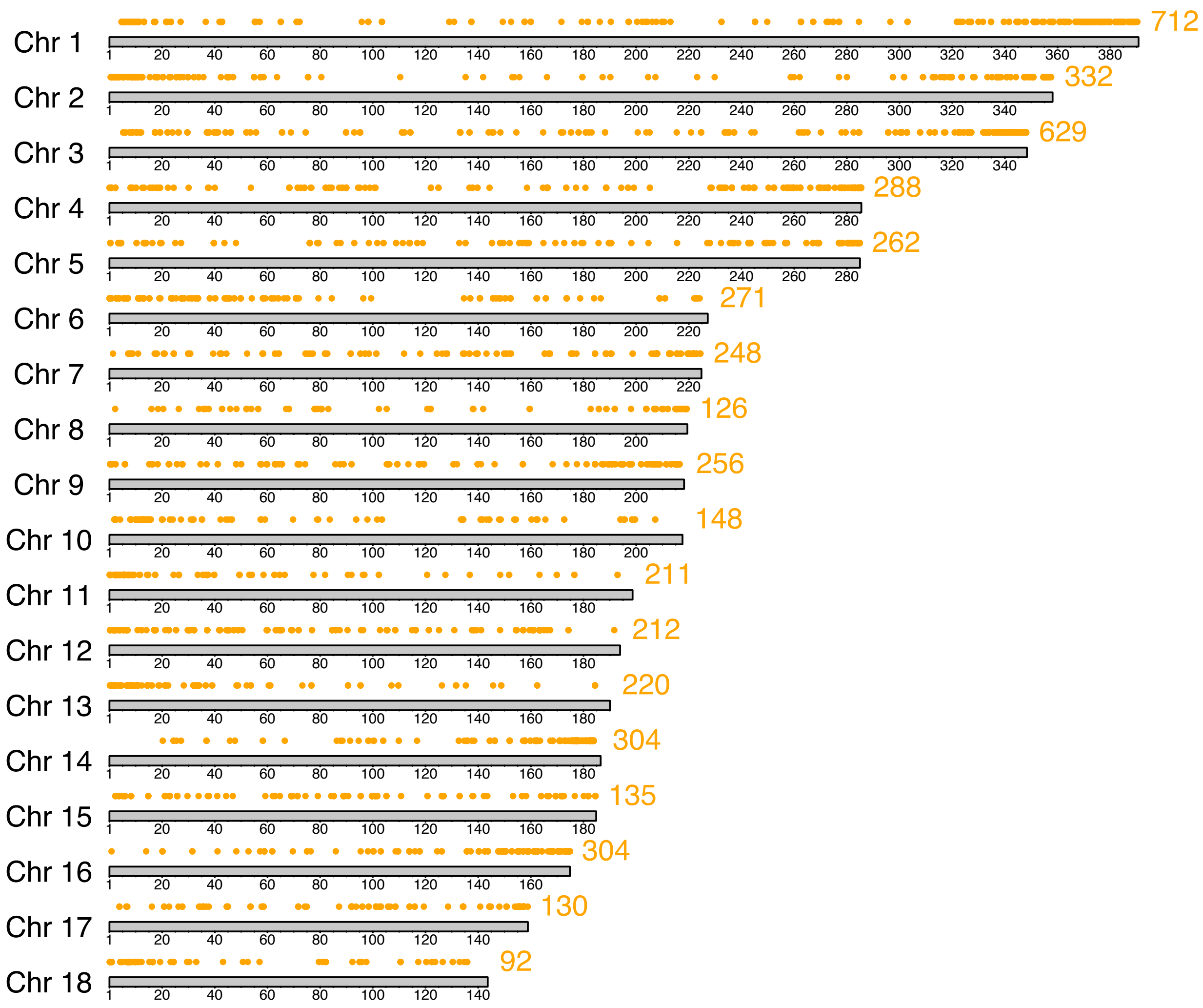

**Fig. S3** Estimates of linkage disequilibrium (LD) over genetic distance for all chromosomes of 86 *Ophrys aveyronensis* individuals. The blue curve indicates the LD decay pattern that was estimated by fitting a trend line based on a nonlinear LOESS regression of r2 on physical distance.

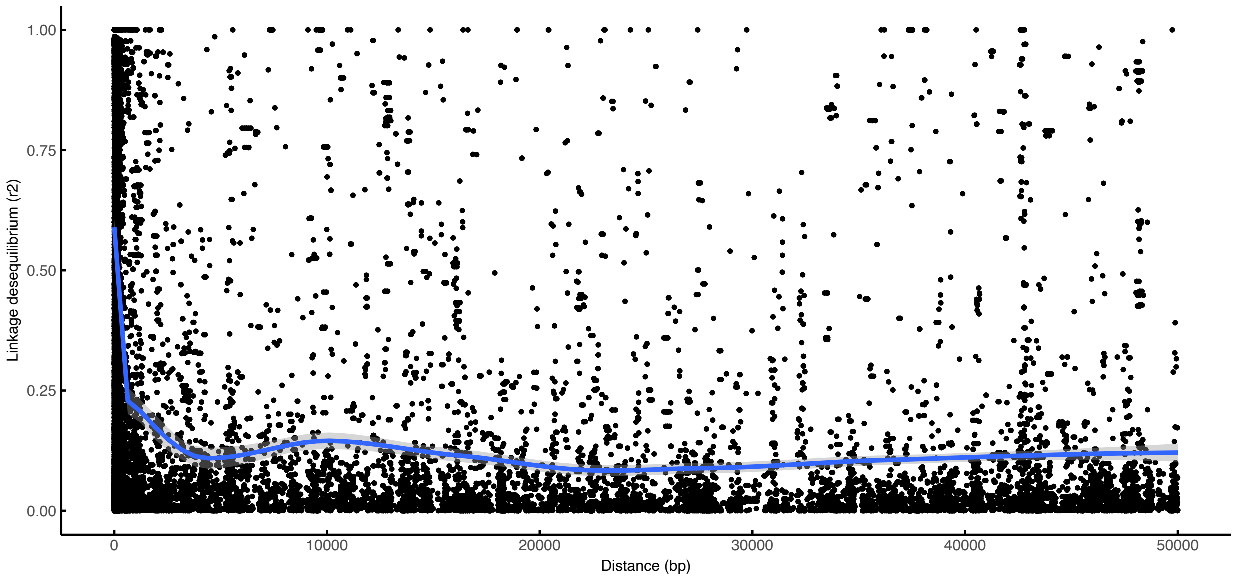


**Fig. S4** Distribution of the correlation coefficient between phenotypic traits (color, odor and morphological).

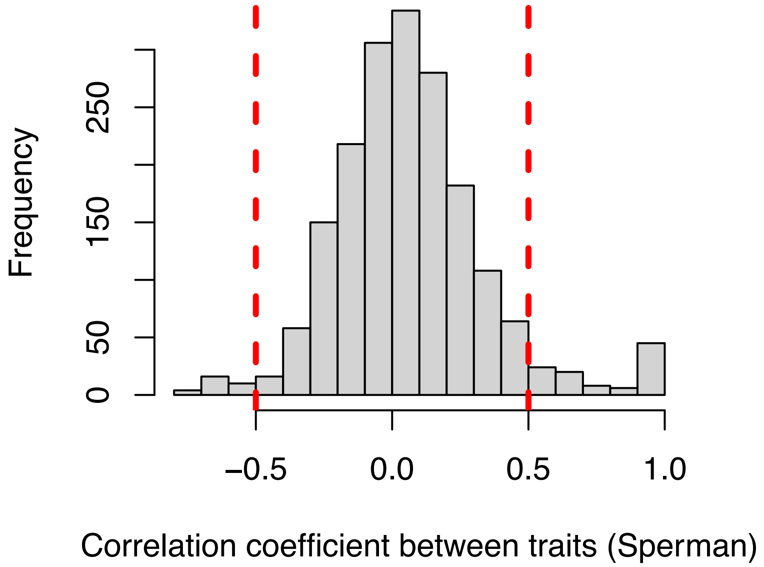

**Fig. S5** Differentiation of floral bouquet between populations with supervised analysis. (**a**) PCA, (**b**) sPLS-DA sample plot with confidence ellipse plots, (**c**) Clustered Image Map (Euclidean Distance) with the 23 odor compounds in column, and the individuals in raw.

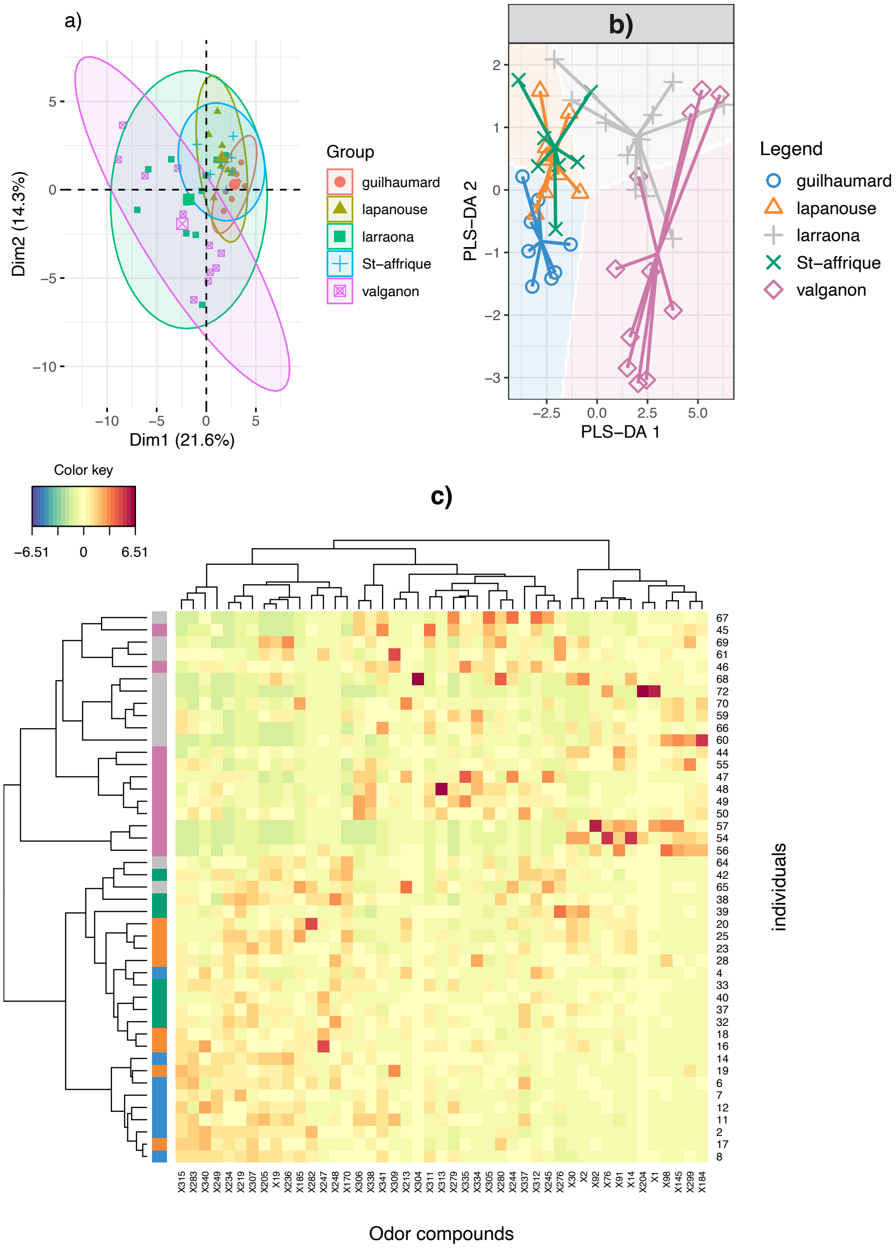

**Fig. S6** Comparisons of phenotypic differentiation ($P_{ST}$) with neutral genetic differentiation ($F_{ST}$) for (**a**) color traits, (**b**) morphological traits, and (**c**) odor traits. $P_{ST}$ (blue line) is plotted with CIs (dotted blue line) as a function of c/h2.

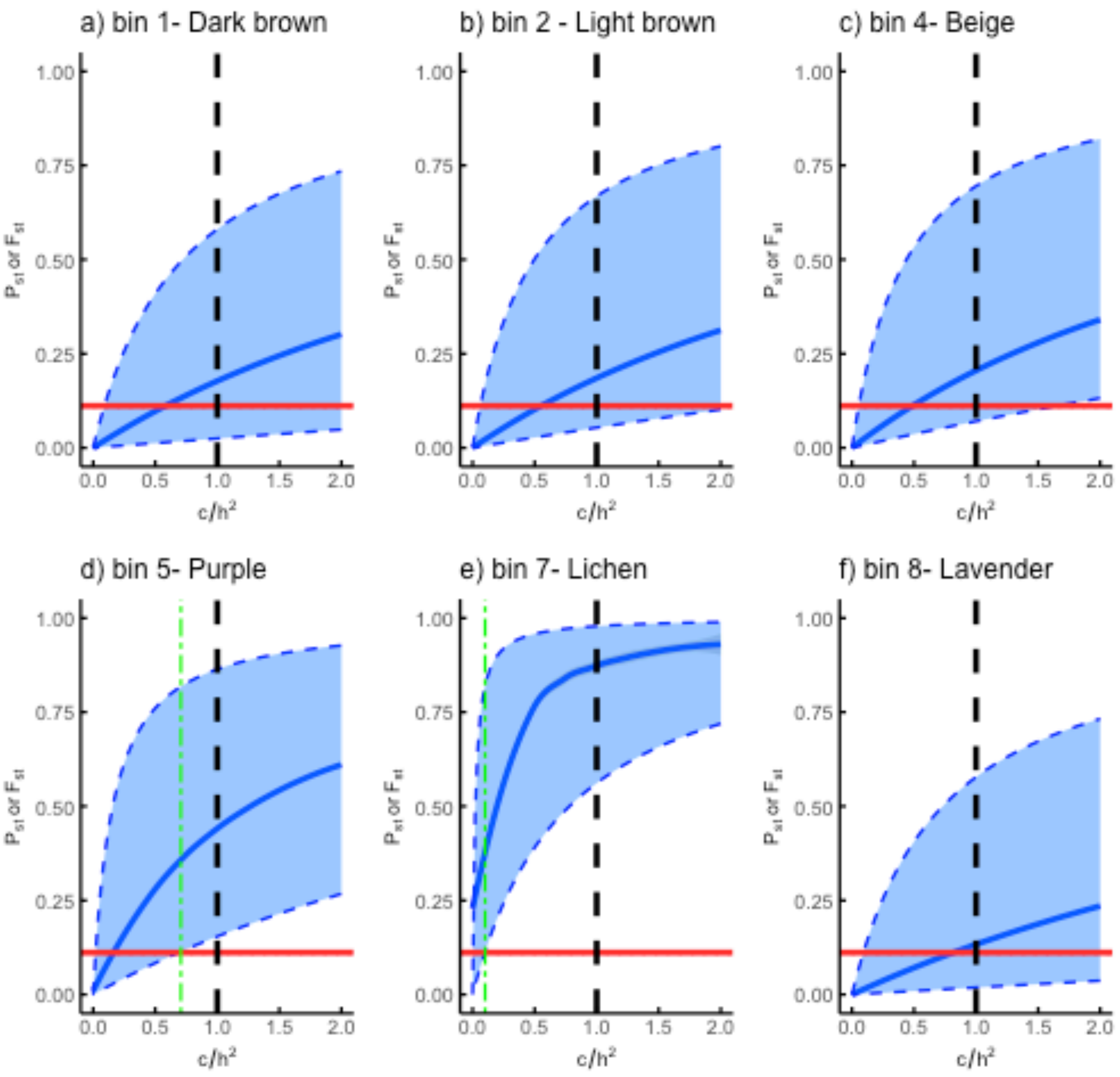

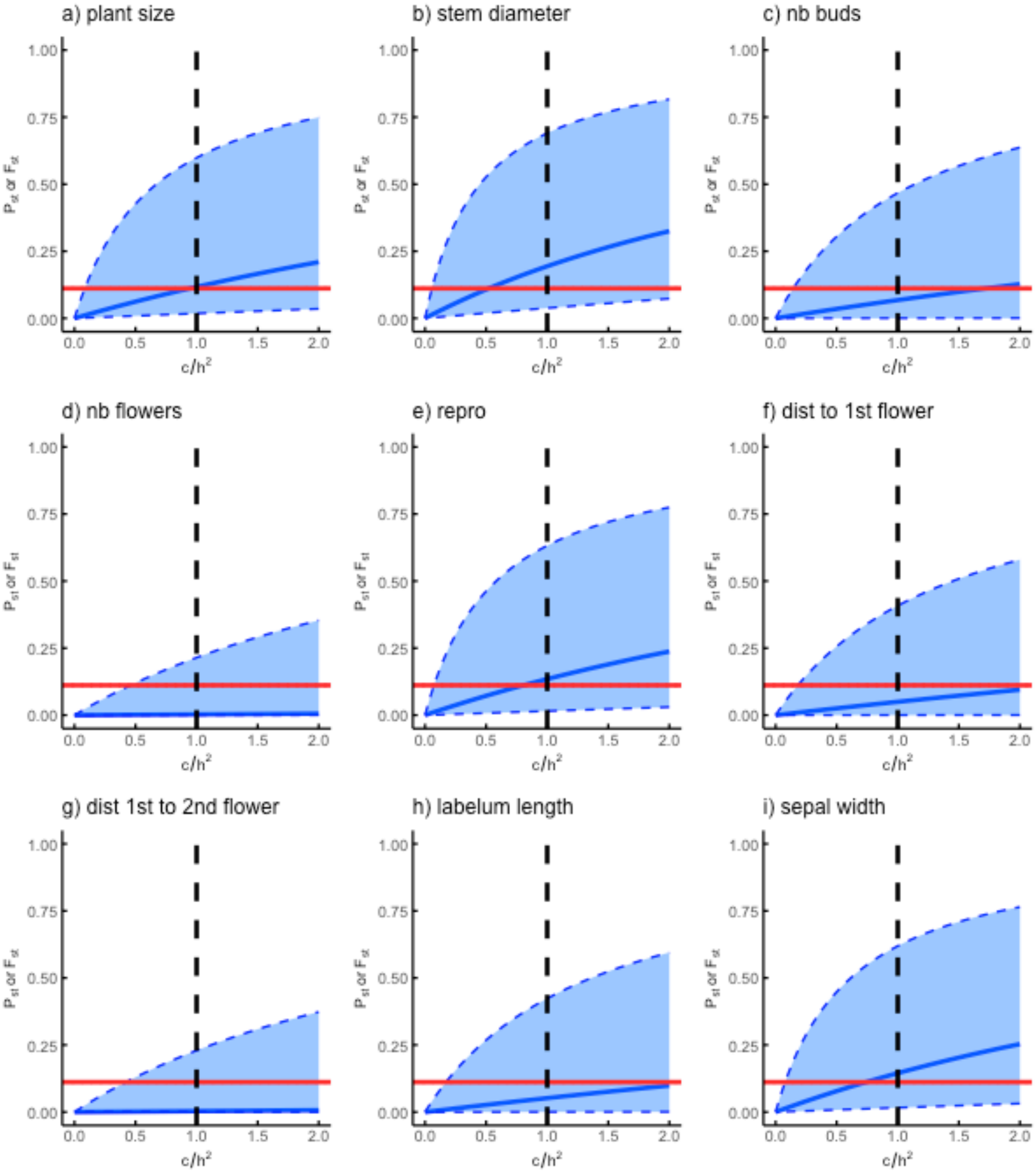

a) plant size
b) stem diameter
c) nb buds
d) nb flowers
e) repro
f) dist to 1st flower
g) dist 1st to 2nd flower
h) labelum length
i) sepal width
$P_{st}$ or $F_{st}$
$c/h^2$

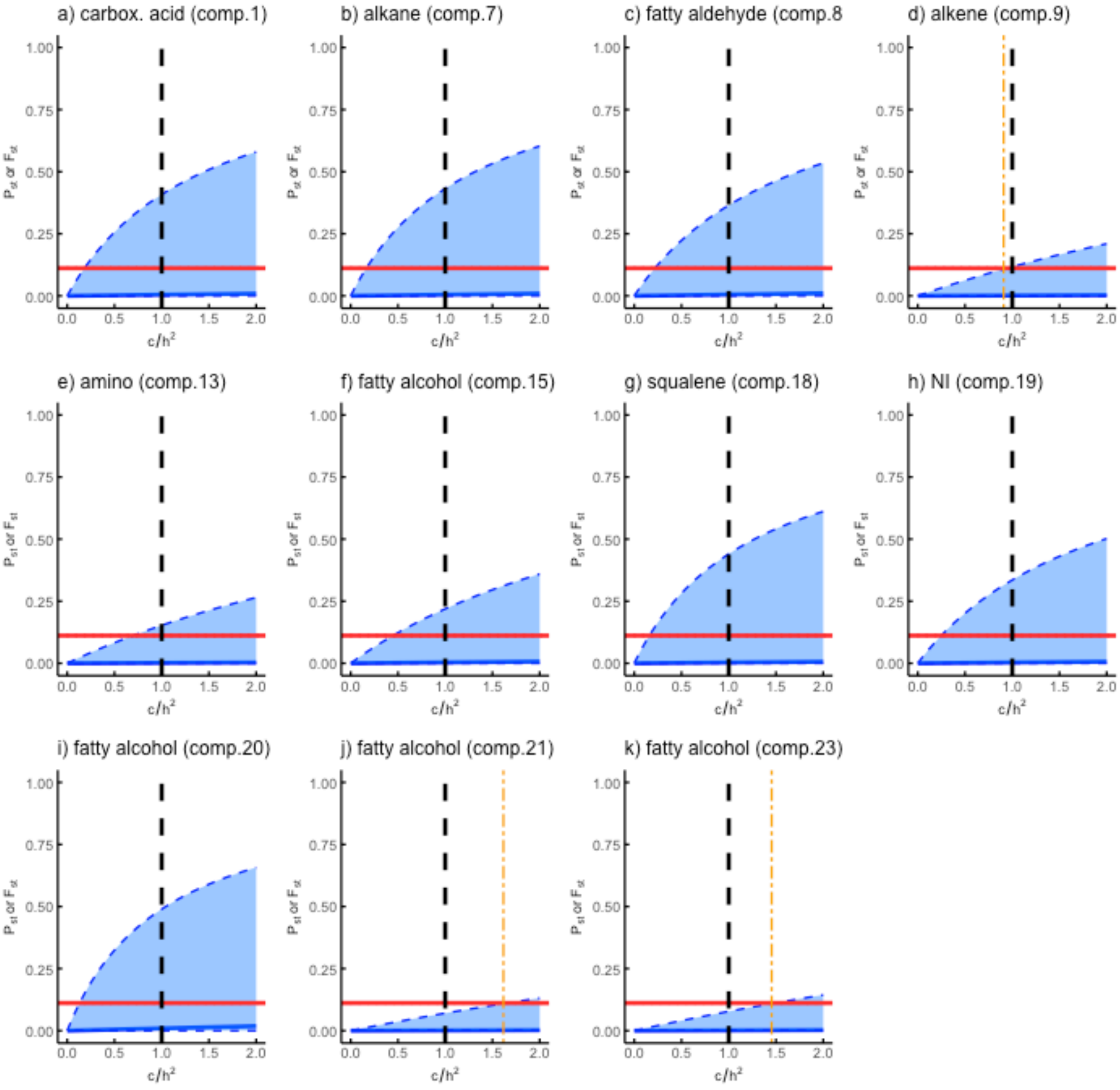
a) carbox. acid (comp.1)
b) alkane (comp.7)
c) fatty aldehyde (comp.8
d) alkene (comp.9)
e) amino (comp.13)
f) fatty alcohol (comp.15)
g) squalene (comp.18)
h) NI (comp.19)
i) fatty alcohol (comp.20)
j) fatty alcohol (comp.21)
k) fatty alcohol (comp.23)
$P_{st}$ or $F_{st}$
$c/h^2$
0.00
0.25
0.50
0.75
1.00
0.0
0.5
1.0
1.5
2.0

**Fig. S7** Manhattan plots depicting outliers SNPs detected by genotype-environment analysis (GEA) on PC1 and PC2 for the *Ophrys aveyronensis* complex. Significant outlier loci are represented by orange dots. Labels refer to the chromosome position of outliers SNP. Red line is the genomic threshold (e.g. -log10(0.01/number of SNPs) and green line the suggestive threshold (e.g. -log10(1/number of SNPs) above which a SNP is significant. Unanchored SNPs are represented in the region beyond chromosome 19. Bio 1: annual mean temperature, Bio 4: temperature seasonality, Bio 5: max temperature of warmest month, Bio 6: min temperature of coldest month, Bio 12: annual precipitation, Bio 13: precipitation of wettest month, Bio 15: precipitation seasonality.

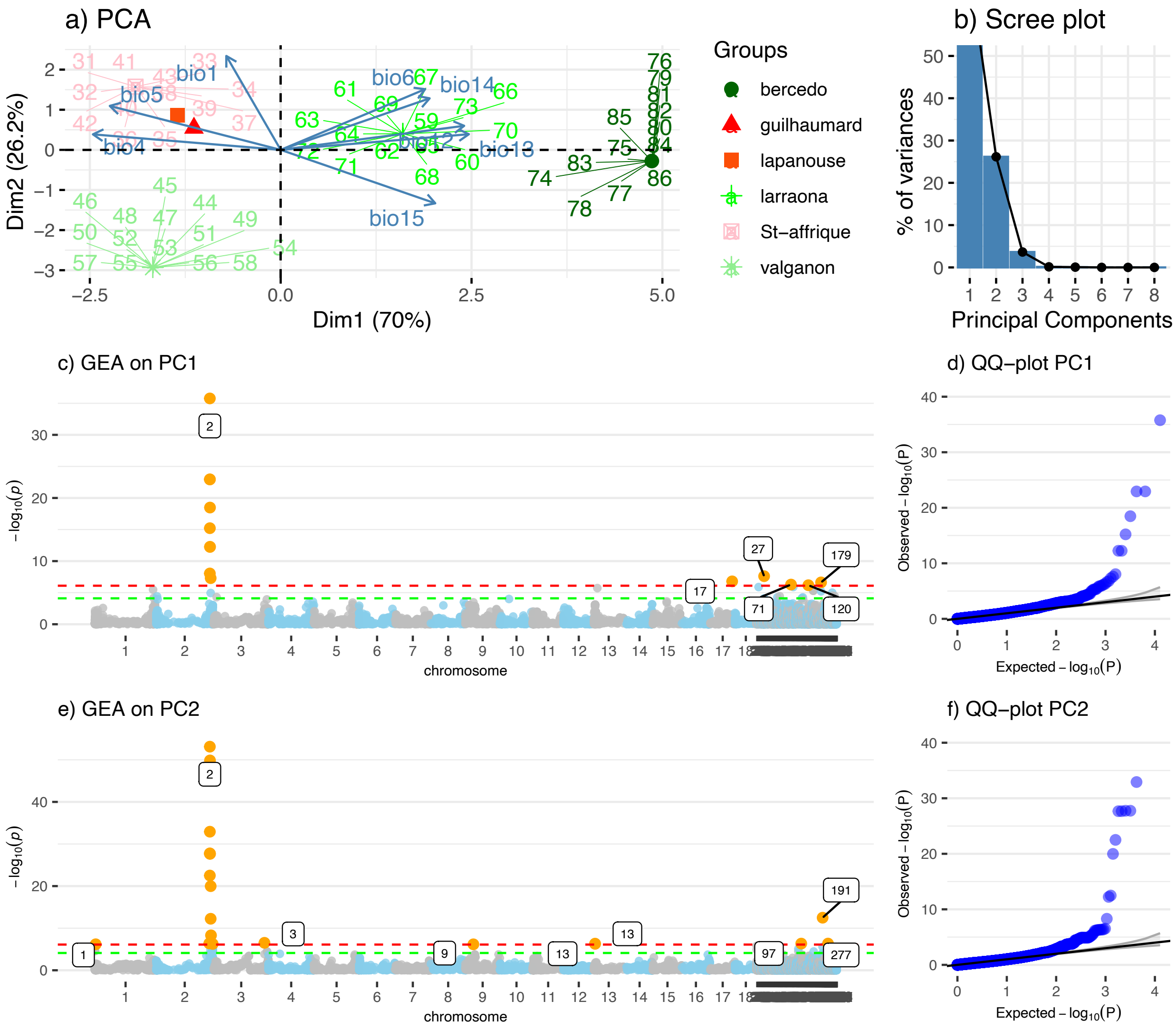

**Fig. S8** Principal component analysis on the odor traits of cluster 1.

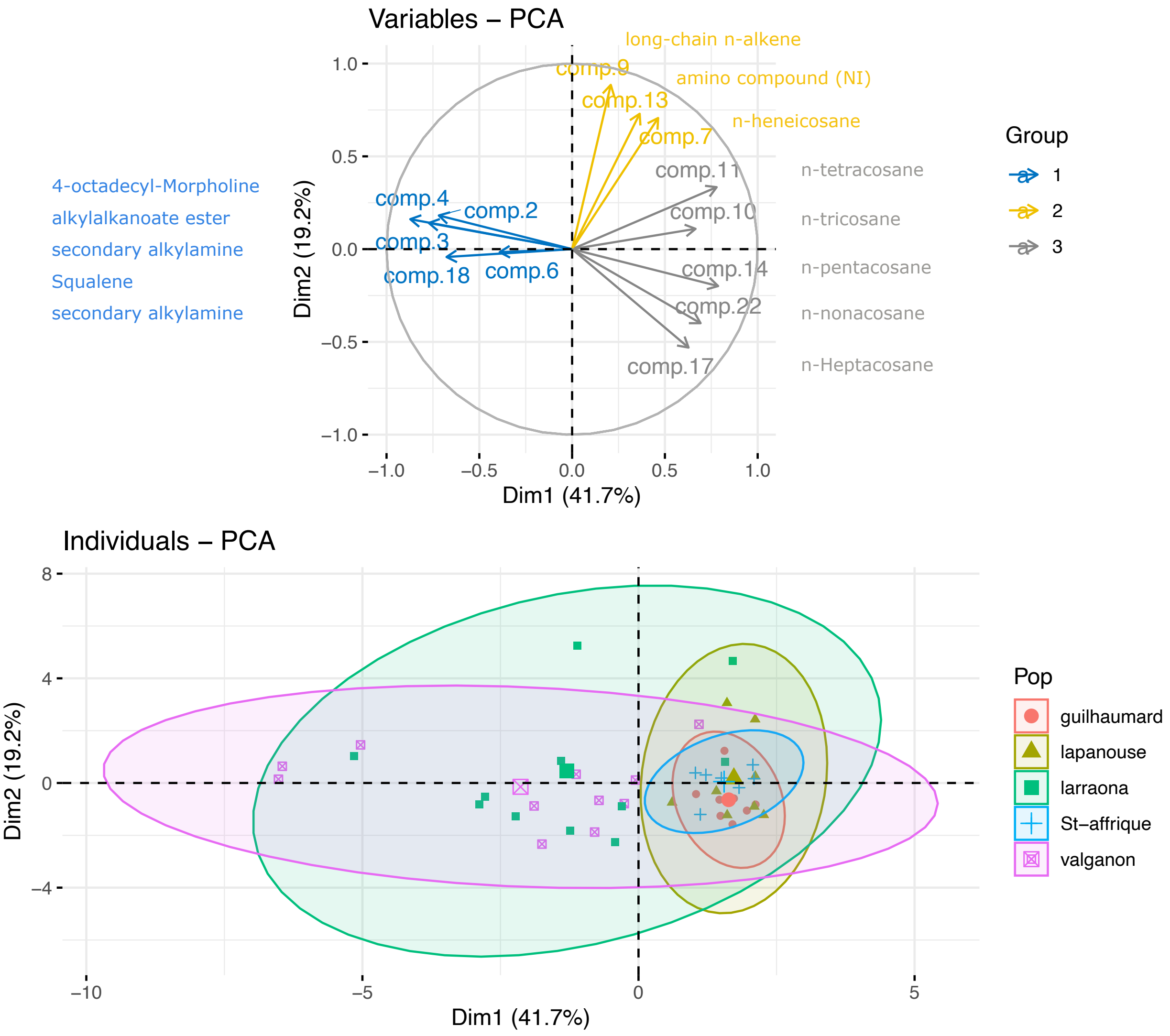

**Fig. S9** Principal component analysis on the color traits.

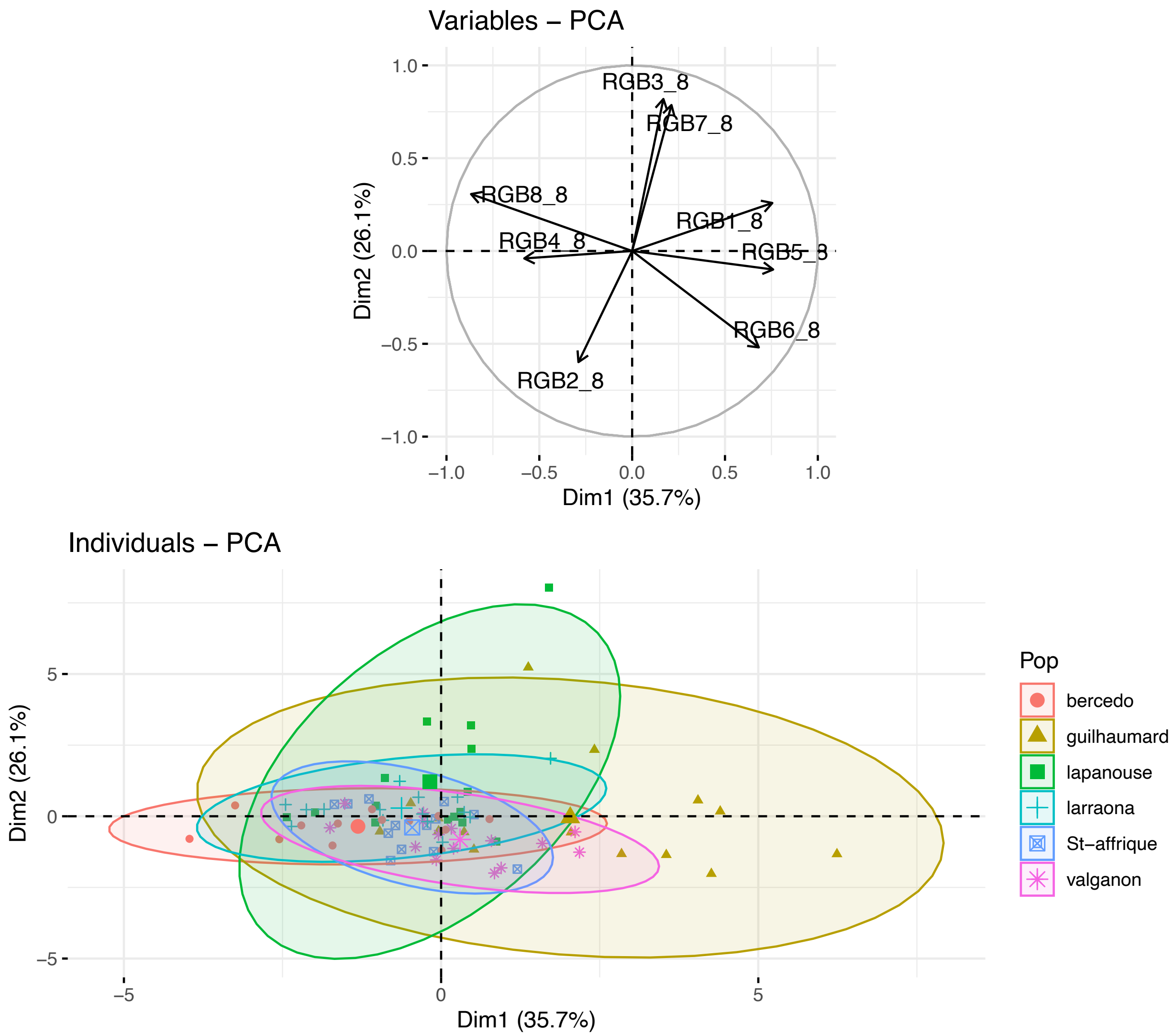


**Methods S1** Acquisition of data on floral volatiles compounds.

The whole surface of the labellum of one unpollinated, intact and fully open flower per individual was rubbed (several times on each part on the labellum surface) with SPME fiber (65-um PDLS-DVB diners, Supelco, Sigma-Aldrich, Bellefonte, PAS, USA) during 1 min and then stored at -20°C before analysis. Flower scents were analyzed by GC-MS using a Shimadzu QP2010 Plus GC-MS with a capillary OPTIMA®5-MS column (30m x 0.25mm, Macherey-Nagel, Düren, Germany) and helium as a carrier gas. SPME fibers were desorbed splitless with an injector temperature of 260 °C and a furnace temperature of 120 °C for 1 min, increased from 120 to 300 °C at a rate of 4 °C/min and held at 300 °C for 2 min. Retention times of a range of n-alkanes (qualitative retention time mix, ASTM, Sigma Aldrich®) were used to convert retention times to retention index. Identification was based on retention index and mass spectroscopy, which were compared with database (NIST 2007, Wiley Registry 9th) and literature (Adams, 2007) and, for some compounds, with analytical standard retention index and mass spectroscopy. Peak areas were measured using GCMS solution software version 4.11 (Shimazu®). The compounds from the control odors are removed from the odor of each flower. A total of 136 compounds were detected in the blends of *O. Aveyronensis.*

**References:** **Adams RP. 2007.** Identification of essential oil components by gas chromatography, quadrupole mass spectroscopy. Carol Stream, IL: Allured Publishing.

**Methods S2** DNA extraction, genotyping and variant calling.

Raw reads were demultiplexed and cleaned with process_radtags, clean reads were mapped onto the reference with bwa v0.7.17 (Li & Durbin, 2009) with default parameters. The loci were built and the SNPs were called thanks to the script ref_map, before final filtering with populations. We kept SNPs that were found in all six populations and in at least 80% of the individuals per population (-r 80), while loci exhibiting an heterozygosity > 70% (--max-obs-het 0.7) were filtered out to reduce the risk of including remaining paralogs loci. We also discarded sites whose minor allele frequency was lower than 5% (--min-maf 0.05). After filtering, 12745 SNPs were kept for subsequent analyses (Fig S2). The LD values between pairs of SNPs in the same chromosome were determined from the squared correlation coefficients (r2) values within the 50-kb window using VCFtools ver. 0.1.16. Pairwise LDs were plotted against physical distances. Physical distances between adjacent markers ranged from 1.00 bp to 50 kb (mean, 0.7 kb). The LD decay pattern was estimated by fitting a trend line based on a nonlinear LOESS regression of r2 on physical distance. This regression curve pattern showed that LD decayed to relatively low levels (r2 < 0.13) within 10 kb (Fig S3). The mean LD between adjacent SNPs was r2 = 0.35. We had a total of 12744 SNPs.

**References:** **Li H, Durbin R. 2009.** Fast and accurate short read alignment with Burrows-Wheeler transform. Bioinformatics. 25:1754-1760. doi: 10.1093/bioinformatics/btp324.

**Methods S3** Patterns of traits variation and integration.

Following He *et al.* (2020), a matrix of trait-trait relationships was first calculated using Spearman correlations. To avoid considering spurious correlations, only significant correlations were considered ($P$ <0.05) and a threshold was applied to determine whether there was a correlation between traits ($|r|$ >0.5) This threshold has been chosen arbitrarily based on the distribution of the correlation coefficients across traits (Fig S4). An agency matrix of a weighted graph was constructed, assigning above the threshold as the coefficient of correlation and below the threshold (or not significant) as 0. Finally a PTN was constructed using the *igraph* package in R software (Csárdi *et al.*, 2023) to visualize the relationships between traits. We used the standard Kamada-Kawai layout, which reflects the relative correlation strengths of the set of nodes in the network. We identified clusters of traits that exhibit covariation among themselves relatively independently of other clusters using the function *cluster_edge_betweenness.* This function identifies modules of traits that are densely connected to each other but sparsely connected to other modules (Newman & Girvan, 2004; Newman, 2006). Substantially similar results were obtained with other community detection algorithms (fast and greedy and multilevel algorithms, results not shown). To test the quality of the partition that has been defined, we used the *modularity* function. This function calculates a modularity index, Q, which is the fraction of edges falling within groups minus the expected fraction of edges in an equivalent network with edges placed as random. The modularity ranges from [-1; 1], with positive values indicating the possible presence of clusters. Traits within a cluster would be studied concomitantly in subsequent GWAS analysis.

**References: Csárdi G, Nepusz T, Müller K, Horvát S, Traag V, Zanini F, Noom D. 2023.** igraph for R: R interface of the igraph library for graph theory and network analysis. **He N, Li Y, Liu C, Xu L, Li M, Zhang J, He J, Tang Z, Han X, Ye Q, et al. 2020.** Plant trait networks: improved resolution of the dimensionality of adaptation. Trends in Ecology & Evolution 35: 908–918. **Newman MEJ. 2006.** Modularity and community structure in networks. Proceedings of the National Academy of Sciences 103: 8577–8582. **Newman MEJ, Girvan M. 2004.** Finding and evaluating community structure in networks. Physical Review E 69: 026113.

**Methods S4** Quantitative genetic approach.

$P_{ST}$ were calculated using to the equation proposed by Brommer (2011):

$$P_{ST} = \frac{\frac{C}{h^2} \cdot \sigma_B^2}{\frac{C}{h^2} \cdot \sigma_B^2 + 2\sigma_W^2}$$

Where $\sigma^2_B$ and $\sigma^2_W$ are the phenotypic variance between and within populations, respectively, the scalar *c* is the proportion of the between-population variance due to genetic effects across populations (rather than phenotypic plasticity), and $h^2$ is the heritability of the trait (i.e. the proportion of phenotypic variance due to additive genetic effects). $P_{ST}$ is easier to measure under common garden conditions/reciprocal transplants, which remove the contribution of environmental effects to phenotypic variation across populations (Luquet et al., 2015). However, such experiments are not feasible for this *Ophrys* species which is both protected and almost impossible to sow or transplant. Therefore, we performed sensitivity analyses as described by Brommer (2011) to quantify the sensitivity of our results to variations in the value of the $c/h^2$ ratio, especially when $c/h^2 < 1$).

To calculate the global $P_{ST}$ for each of the phenotypic traits, we extracted the variance components ($\sigma^2_B$ and $\sigma^2_W$) and their confidence intervals (CIs) from the posterior distribution of a Bayesian generalized linear mixed model (Bertrand *et al.*, 2016; Seeholzer & Brumfield, 2018). This approach provides more accurate and less biased estimates of $P_{ST}$ than other methods (O'Hara & Merilä, 2005; Leinonen *et al.*, 2008). We combined populations of both subspecies to perform $P_{ST}$–$F_{ST}$ comparison in order to obtain reasonable estimates of $P_{ST}$ (O'Hara & Merilä, 2005). Bayesian generalized mixed models were performed in R using the *MCMCglmm* package (Hadfield, 2010), with a weakly informative Inverse-Gamma distribution for priors. Where necessary, traits were log-transformed to meet model assumptions. We excluded from this analysis, traits for which models did not converge and/or did not meet model assumptions (Table S1, Criterion 3).

References: **Bertrand JAM, Delahaie B, Bourgeois YXC, Duval T, García-Jiménez R, Cornuault J, Pujol B, Thébaud C, Milá B. 2016.** The role of selection and historical factors in driving population differentiation along an elevational gradient in an island bird. Journal of Evolutionary

Biology 29: 824–836. **Brommer JE. 2011.** Whither Pst? The approximation of Qst by Pst in evolutionary and conservation biology: Whither Pst? Journal of Evolutionary Biology 24: 1160–1168. **Leinonen T, O'Hara RB, Cano JM, Merilä J. 2008.** Comparative studies of quantitative trait and neutral marker divergence: a meta-analysis: QST - FST meta-analysis. Journal of Evolutionary Biology 21: 1–17. **Luquet E, Léna J-P, Miaud C, Plénet S. 2015.** Phenotypic divergence of the common toad (Bufo bufo) along an altitudinal gradient: evidence for local adaptation. Heredity 114: 69–79. **O'Hara RB, Merilä J. 2005.** Bias and Precision in Q ST Estimates: Problems and Some Solutions. Genetics 171: 1331–1339. **Hadfield JD. 2010.** MCMC Methods for Multi-Response Generalized Linear Mixed Models: The MCMCglmm R Package. Journal of Statistical Software 33. **Seeholzer GF, Brumfield RT. 2018.** Isolation by distance, not incipient ecological speciation, explains genetic differentiation in an Andean songbird (Aves: Furnariidae: *Cranioleuca antisiensis*, Line-cheeked Spinetail) despite near threefold body size change across an environmental gradient. Molecular Ecology 27: 279–296.

**Methods S5** Forward genetic approaches.

*Pcadapt* is an outlier detection method based on principal component analysis. This method detects population structure from genetic data using a PCA, and then calculates statistics and p-values for the association between each SNP and the first K principal components (PCs). We first assess the optimal K value from 1 to 10 using a screen plot of the proportion of variance explained by each PC. We kept K= 5 which is also the optimal number of genetic groups found in previous results (Gibert *et al.,* 2023). To identify outliers associated with subspecies divergence, we also ran the function with K=2 corresponding to each subspecies. We then calculated q-value (i.e. p-values adjusted for multiple comparisons) using the R package "qvalue" (Storey *et al.,* 2015). Finally, we obtained a list of significant SNPs with an expected false discovery rate (FDR) of 0.01, meaning that 1% of the candidate SNPs are expected to be false positives. The analysis was performed using the version 4.3.3 of the R package "PCAdapt" (Luu *et al.,* 2017; Privé *et al.* 2020).

The *sNMF* method uses the sparse non-negative matrix factorization algorithm implemented in the R package LEA (*snmf* function, Frichot & François 2015) to compute estimates of ancestry coefficients, and ancestral allele frequencies. Genome scans for adaptive alleles were then performed based on population differentiation statistics computed from the ancestral allele frequencies (*snmf_pvalue* function). Here, we tested the number of genetic clusters k= 5 and k=2 to test for population and subspecies adaptive divergence. Before running the *snmf* function, we imputed missing genotypes based on the estimated ancestry coefficient and on ancestral genotype frequencies using the function *impute*. To obtain a list of candidate SNPs, we transformed p-values to q-values and applied the same filtering criteria as in *Pcadapt*.

*BayPass* (Bayesian population association analysis, Gauthier, 2015) is a command-line executable program that uses Bayesian hierarchical models. Divergence at each locus was characterized using the XtX statistic, a measure of adaptive differentiation corrected for population structure and demography (Günther & Coop, 2013). *BayPass* was run with default parameters under the core model: MCMC chains were run for 25000 iterations after a burn-in period of 5000 iterations. Here, we computed contrasts of standardized allele frequencies between two groups of populations (France vs Spain) across the 6 populations. For each SNP, *BayPass* computed XtX statistics and its corresponding p-value (on a -log10 scale), assuming a Chi2 distribution. To obtain a list of candidate SNPs, we transformed p-values to q-values and applied the same filtering criteria as *Pcadapt* and *sNMF.*

*GEA:* Parameters with the *lfmm2* function were estimated using a frequentist approach with least squares estimation (recommended for large genotype matrices > 10.000 SNPs). Missing genotypes were replaced by imputed values using a missing data imputation method provided in the LEA package (function *impute)*. To obtain a list of candidate SNPs, we transformed p-values to q-values and applied an expected false discovery rate (FDR) of 0.01, meaning that only 1% of the candidate SNPs are expected to be false positives

**References:** **Frichot E, François O. 2015.** LEA: An R package for landscape and ecological association studies (B O'Meara, Ed.). Methods in Ecology and Evolution 6: 925–929. **Gautier M. 2015.** Genome-Wide Scan for Adaptive Divergence and Association with Population-Specific Covariates. Genetics. 201(4):1555-79. doi: 10.1534/genetics.115.181453. **Gibert A, Buscail R, Baguette M, Fraïsse C, Roux C, Schatz B, Bertrand JAM. 2023.** Climate change promoted allopatric divergence and explains the current disjunct geographic distribution of the Ophrys aveyronensis species complex (Orchidaceae): 2023.04.27.538532. **Günther T., Coop G., 2013.** Robust identification of local adaptation from allele frequencies. Genetics 195: 205–220. **Luu K,**

**Bazin E, Blum MG. 2017.** pcadapt: an R package to perform genome scans for selection based on principal component analysis. *Mol Ecol Resour*. 17(1):67–77. **Privé F, Luu K, Vilhjálmsson BJ, Blum MGB. 2020.** Performing Highly Efficient Genome Scans for Local Adaptation with R Package pcadapt Version 4, *Molecular Biology and Evolution*, 37: 2153–2154_**Storey JD, Bass AJ, Dabney A, Robinson D. 2015.** qvalue: Q-value estimation for false discovery rate control R package version 2120.

**Methods S6** Reverse genetic approaches.

For both FarmCPU and BLINK, the first three principal components were fitted as covariate variables to reduce the false positives due to population structure. In addition, they incorporate associated markers as covariate (multi-locus models) to test the marker and eliminate their connection to the cryptic relationship among individuals (kinship effect). For BLINK, the associate markers are selected according to linkage disequilibrium optimized for Bayesian information content and reexamined across multiple tests to reduce false negatives. In terms of statistical power and computational efficiency, simulation studies have shown that BLINK is superior to FarmCPU, both being largely superior to linear models or general linear models (Huang *et al.* 2019).

**References:** **Huang M, Liu X, Zhou Y, Summers RM, Zhang Z. 2019.** BLINK: a package for the next level of genome-wide association studies with both individuals and markers in the millions. GigaScience 8: giy154.

**Table S1** Criteria for inclusion of traits in the analysis.

**Table S2** Phenotypic differentiation among populations and regions for 45 traits in *Ophrys aveyronensis*. Results from GLMs. Significant results are in bold. ns = non significant; ***P < 0.001, **P < 0.01, *P < 0.05. Several odor traits were also tested as binomial variables.

**Table S3** List of SNPs identified by several GWAS methods by trait. SNP: ID, CHR: chromosome, BP: position in base pairs of the SNP found by several methods. Sum: number of methods which found this SNP. Chrs >18 refers to unanchored regions.

**Table S4** List of SNPs identified by GWAS methods and located within or close to an annotated CDS CHRS: chromosome, BP: position in base pairs of the SNP. SumGWAS: number of GWAS models which found this SNP. Chrs >18 refers to unanchored regions. Up: start position of the CDS in bp, Down : end position of the CDS in bp. length: length of the sequence (bp), match: length of match on reference (bp), pvalue for GWAS run with BLINK, lfmm2 and FarmCPU models.

**Table S5** GO enrichment analysis on region of 7Mb of chromosome 2.

**Table S6** Eggnog annotation of genes on 7Mb of chromosome 2.

**Table S7** Eggnog annotation of genes on 22Mb of chromosome 2.
